# Uncovering the Infection Strategy of *Phyllachora maydis* during Maize Colonization: A Comprehensive Analysis

**DOI:** 10.1101/2023.08.26.554799

**Authors:** Denise L. Caldwell, Camila R. Da Silva, Austin G. McCoy, Harryson Avila, John C. Bonkowski, Martin I. Chilvers, Matthew Helm, Darcy E. Telenko, Anjali S. Iyer-Pascuzzi

## Abstract

Tar spot, a disease caused by the ascomycete fungal pathogen *Phyllachora maydis*, is considered one of the most significant yield-limiting diseases of maize (*Zea mays* L.) within the United States. *P. maydis* may also be found in association with other fungi, forming a disease complex with characteristic fish eye lesions. Understanding how *P. maydis* colonizes maize leaf cells is essential for developing effective disease control strategies. Here, we used histological approaches to elucidate how *P. maydis* infects and multiplies within susceptible maize leaves. We collected tar spot-infected maize leaf samples from four different fields in northern Indiana at three different time points during the growing season. Samples were chemically fixed and paraffin-embedded for high-resolution light and scanning electron microscopy. We observed a consistent pattern of disease progression in independent leaf samples collected across different geographical regions. Each stromata contained a central pycnidium that produced asexual spores. Perithecia with sexual spores developed in the stomatal chambers adjacent to the pycnidia, and a cap of spores formed over the stromata. *P. maydis* reproductive structures formed around but not within the vasculature. In our samples containing fish eye lesions, *P. maydis* is associated with two additional fungi, one of which is likely a member of the *Paraphaeospheria* genus; the other is an unknown fungi. Our data provide fundamental insights into how this pathogen colonizes and spreads within maize leaves. This knowledge can inform new approaches to managing tar spot, which could help mitigate the significant economic losses caused by this disease.

## INTRODUCTION

Plant diseases caused by fungi are among the most severe limiting factors in global crop production (Almeida et al. 2019; Stukenbrock and Gurr 2023). Such diseases cause an estimated 10-20% yield loss worldwide annually, with an additional 10%-20% loss post-harvest (Stukenbrock and Gurr 2023). *Phyllachora maydis* is an obligate fungal pathogen that causes tar spot, a foliar disease of maize (*Zea mays* L.) common in the United States, Mexico, Central and South America (Bajet et al. 1994; Mottaleb et al. 2019; Rocco da Silva et al. 2021; Solórzano et al. 2023; Valle-Torres et al. 2020). Tar spot was first detected in the U.S. in 2015 (Collins et al. 2021; Malvick et al. 2020; Mottaleb et al. 2019; Moura et al. 2023; Pandey et al. 2022; Rocco da Silva et al. 2021; Ruhl et al. 2016; Valle-Torres et al. 2020; Wise et al. 2023). Since then, *P. maydis* has caused an estimated $2.9 billion in maize yield loss (Mueller et al. 2021, 2022). *P. maydis* produces dark-pigmented structures known as stromata, the characteristic “tar spots” visible on the leaf surface (Rocco da Silva et al. 2021; Solórzano et al. 2023; Valle-Torres et al. 2020). Later in the disease cycle, a necrotic halo may develop surrounding the stromata, giving the appearance of a “fish eye” (Hock et al. 1995; McCoy et al. 2019; Mottaleb et al. 2019; Rocco da Silva et al. 2021; Valle-Torres et al. 2020). Historically in Central and South America, there have been three fungal species reported in the tar spot disease complex; *P. maydis*, *Monographella maydis* (syn= *Michrodochium maydis*), and *Coniothyrium maydis* (Hock et al. 1992, 1995). *P. maydis* causes tar spot disease through its production of stromata, *C. maydis* was described as the hyperparasite of *P. maydis*, and *M. maydis* was believed to cause the necrotic halo around *P. maydis* termed the fish eye lesion. However, *M. maydis* has not been associated with fish eye lesions in the U.S. (McCoy et al. 2019), and whether *C. maydis* is part of the tar spot disease complex present in the U.S. is unclear (Rocco da Silva et al. 2021).

Detecting the early infection of *P. maydis* can be challenging, as signs of the disease may not be present for 2 – 3 weeks after fungal infection (Kleczewski et al. 2019; Solórzano et al. 2023). Thus, tar spot may go unnoticed until *P. maydis* has colonized the underlying leaf cells. Genetic resistance is not well described (Cao et al. 2017, 2021; Ceballos 1992; Mahuku et al. 2016; Yan et al. 2022) and disease control centers primarily on fungicide applications (Rocco da Silva et al. 2021; Telenko et al. 2020, 2022a, 2022b; Valle-Torres et al. 2020). Disease management strategies are further hampered by our limited understanding with regards to how *P. maydis* invades, colonizes and spreads within maize leaf cells and across leaves. Previous work examined colonization of *P. ischaemi* and *P. parilis* in the grass species *Ischaemum australe* and *Paspalurn orbiculare* respectively (Parbery 1963b), but microscopic studies examining the cellular infection of *P. maydis* have not been reported. Whether colonization and spread of *P. maydis* in maize leaves is similar to that of *P. ischaemi* and *P. parilis* is unknown but is important knowledge for developing improved disease control strategies.

Here we sought to further elucidate the infection strategy used by *P. maydis* to colonize maize. We sampled leaves from susceptible hybrid corn at three different time points throughout the growing season in the U.S. state of Indiana. Using histological-based approaches coupled with light and scanning electron microscopy we investigated how *P. maydis* grows within and emerges from a maize leaf. In addition, we found that the typical fish eye symptom consists of *P. maydis* and at least two additional spore types, a *Paraphaeosphaeria* spp. and an unidentified species. Collectively, the data presented provide the first high-resolution, detailed analysis of *P. maydis* cellular colonization and spread within maize and significantly advances our understanding of the infection strategy used by *P. maydis*.

## MATERIALS & METHODS

### Field Collection

Tar spot infected leaves were collected at the Pinney Purdue Agricultural Center (PPAC) in Porter County, IN (41°41’49.07"N, -86°79’92.49"W), and on-farm sites in St. Joseph County, IN (41°58’63.89"N, -86°43’32.22"W), Porter County, IN (41°46’34.94"N, - 87°00’22.98"W), and LaPorte County, IN (41°41’49.07"N, -86°79’92.49"W). Foliar samples were collected at early, mid, and late disease onset during the 2020 field season. Stage of corn development was recorded during each sampling dating according to (Nleya 2019). As consistent inoculation procedures are not available for studies of *P. maydis* infection in a controlled environment, we relied on field-collected samples; thus, penetration of the appressorium into the epidermis was not observed in conjunction with stromata formation.

Samples were collected at different stages of the disease cycle in maize plants at the blister (R2), milk (R3), dent (R5) and physiological maturity (R6) growth stages (Nleya 2019). On 28 July 2020, samples were collected from plants at R2 growth stage in LaPorte and St. Joseph counties and at R3 growth stage in Porter County. All samples had early signs of tar spot. A second set of leaves was collected on 2 September in LaPorte, St. Joseph, and Porter counties and on 16 September at PPAC, all at the R5 corn growth stage. Leaves showing severe tar spot disease symptoms were collected on 16 September in LaPorte, St. Joseph, and Porter counties, on 23 and 29 September, and on 6 October at PPAC. All were at R6 growth stage.

Six maize leaves with tar spot symptoms and six without symptoms (control) were collected from twelve random plants per field at each time point. At later time points it was not possible to collect leaves without symptoms and thus only the six symptomatic leaves were collected. All leaves were assessed for severity as percent tar spot stroma (0 to 100%) (black spots) following a standardized rating scale developed by Telenko et al. (2020). From each sampled leaf, six tar spot lesions (0.5 cm x 0.5 cm) from the top, middle, or bottom of the leaf were excised using a sterilized scalpel. All lesions were transferred directly into a vial with fixative solution as described below. A total of six vials were collected for each location with six stromata tissue inside each, as well as six vials for each location with six asymptomatic tissues inside each vial. Samples with asymptomatic leaves were not collected in Porter County on 2 September and 16 September as well as in St. Joseph County on 16 September due to lack of leaves without disease symptoms. The vials were placed in a cooler with ice.

### Histology and Microscopy

The 0.5 cm x 0.5 cm samples were collected from the field and placed directly into fixative solution containing 1.5% glutaraldehyde and 2% paraformaldehyde in 0.1 M cacodylate buffer (pH 6.8) on ice. The samples were returned to the laboratory and prepared as described by Caldwell and Iyer-Pascuzzi (2019). Briefly, samples were paraffin-embedded and sectioned in three different orientations: each leaf sample contained six stromata; two were mounted into paraffin wax in the transverse orientation, two in the longitudinal orientation, and two in the lateral orientation. The samples were cut at 12 µm thickness for light microscopy and 20 µm for scanning electron microscopy (SEM). The light sections were stained with 0.05% toluidine blue (Sigma-Aldrich, Inc., U.S.) and visualized on an Olympus BX43 light microscope using a Spot Idea 5.0 Mp Color Digital CMOS camera. Toluidine Blue is a polychromatic dye that preferentially stains acidic tissues and stains multiple tissues within sections (O’Brien et al. 1964). SEM samples were prepared using the same method as previously described (Caldwell and Iyer-Pascuzzi 2019).

### DNA extraction, sequencing and analysis

DNA was isolated using a Quick-DNA Fungal Miniprep Kit (Zymo). DNA was fragmented using a S2 sonicator (Covaris) followed by library construction using an xGen DNA Library Prep MC UNI kit (Integrated DNA Technologies, Inc., U.S.) and indexed using xGen UDI-UMI Adapter (Integrated DNA Technology, Inc., U.S.). Sequence was generated from an NovaSeq 6000 using 300 cycle chemistry to produce two 150 base pair reads. Raw paired-end reads were checked for quality using FastQC, and bases with less than Q20 were trimmed using BBtools (V38.18) (Andrews 2010; Bushnell 2014). Trimmed paired-end reads were then aligned to the B73 maize genome (version 5.0) and the *Phyllachora maydis* PM02 genome using Bowtie2, and unmapped read pairs were used for further analysis (Langmead and Salzberg 2012). DIAMOND (version 2.1.4) and MEGAN (version 6.24.20) BLAST were used to identify fungal reads and classify the unmapped read pairs for each sample to the genus level using the NCBI-nr database (Buchfink et al. 2021; Huson et al. 2016). Genera which had less than 1,000 reads mapped were removed. MEGAN genera abundance results were then exported as text files and visualized in R (V4.1.1) using “ggplot” (R Core Team 2021.; Wickham 2009).

### Data Availability

Raw sequences were submitted to the NCBI Sequence Read Archives. All bioinformatic and analysis code are available here: https://github.com/AGmccoy/Tarspot-histology-microbiome-investigation.

## RESULTS

### Mature *P. maydis* stroma structure consists of four sub-structures: vegetative hyphae, clypeus, pycnidium, and perithecia.

A detailed examination of histological sections (see Supplementary Fig. 1 for definitions of orientation) revealed that an individual stromata, often referred to as a tar spot, is comprised of multiple fungal sub-structures, each displaying a distinct cellular arrangement (Fig. 1). The innermost layer of the stromata houses the fungal reproductive structures, specifically the pycnidium, which is responsible for producing asexual spores (Fig. 1). As the stromata matures, perithecia-generating sexual spores develop around the multilobed pycnidia (Fig. 1; see Supplementary Fig. 2-4 for serial sections through stromata at different disease stages). The outermost layer, the clypeus, likely serves as a protective structure for perithecia and pycnidium (Fig. 1). Vegetative hyphae are consistently found connecting these sub-structures. Collectively, these sub-structures compose a tar spot stromata (Fig. 1).

**Fig. 1.**
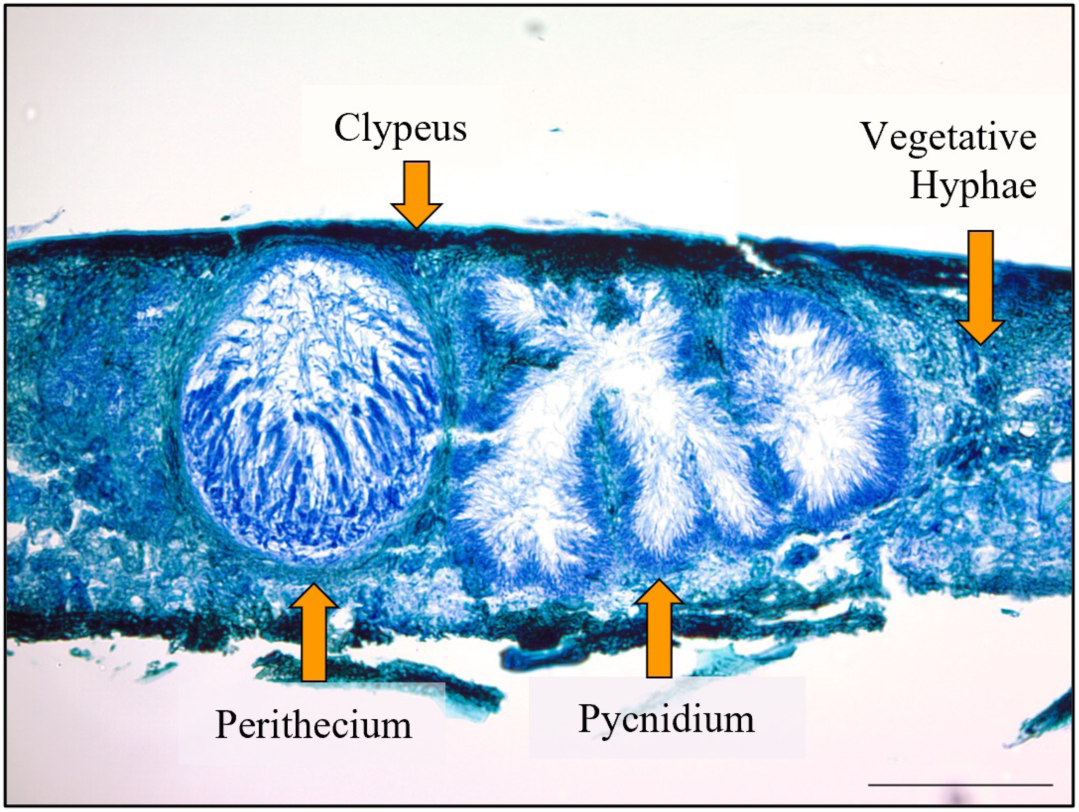
Mature stromata anatomy of *P. maydis*. The mature *P. maydis* stromata is comprised of four different sub-structures. 12 µm section of a maize leaf stained with toluidine blue and viewed with light microscopy. The clypeus is the remnants of the *Z. mays* epidermis filled with densely packed melanized hyphae. Also present are vegetative hyphae, a perithecium which produces sexual spores, and a central multilobed pycnidium which contains asexual spores that exude through a central ostiole (not pictured). Scale bar represents 10 µm.

Members of *Phyllachora* generally exhibit a consistent infection strategy wherein during the initial stages of colonization, ascospores develop a germination tube on the surface of the maize leaf (Parbery 1963b, 1967). The germination tube then forms an appressorium that penetrates the maize epidermis. Although we were unable to observe these initial infection stages, subsequent *P. maydis* development appears similar to that of other *Phyllachora* species. Hyphae fill the epidermal cells, creating the clypeus. A central asexual structure called the pycnidium forms (Fig. 2A). As the stroma continues to mature, multiple sexual perithecia develop near the pycnidium (Fig. 2B-D). Throughout the progression of the disease, the central pycnidium locule becomes increasingly lobed, while the ascomata within the stroma mature into fully developed sexual structures (Fig. 2B-D).

**Fig. 2.**
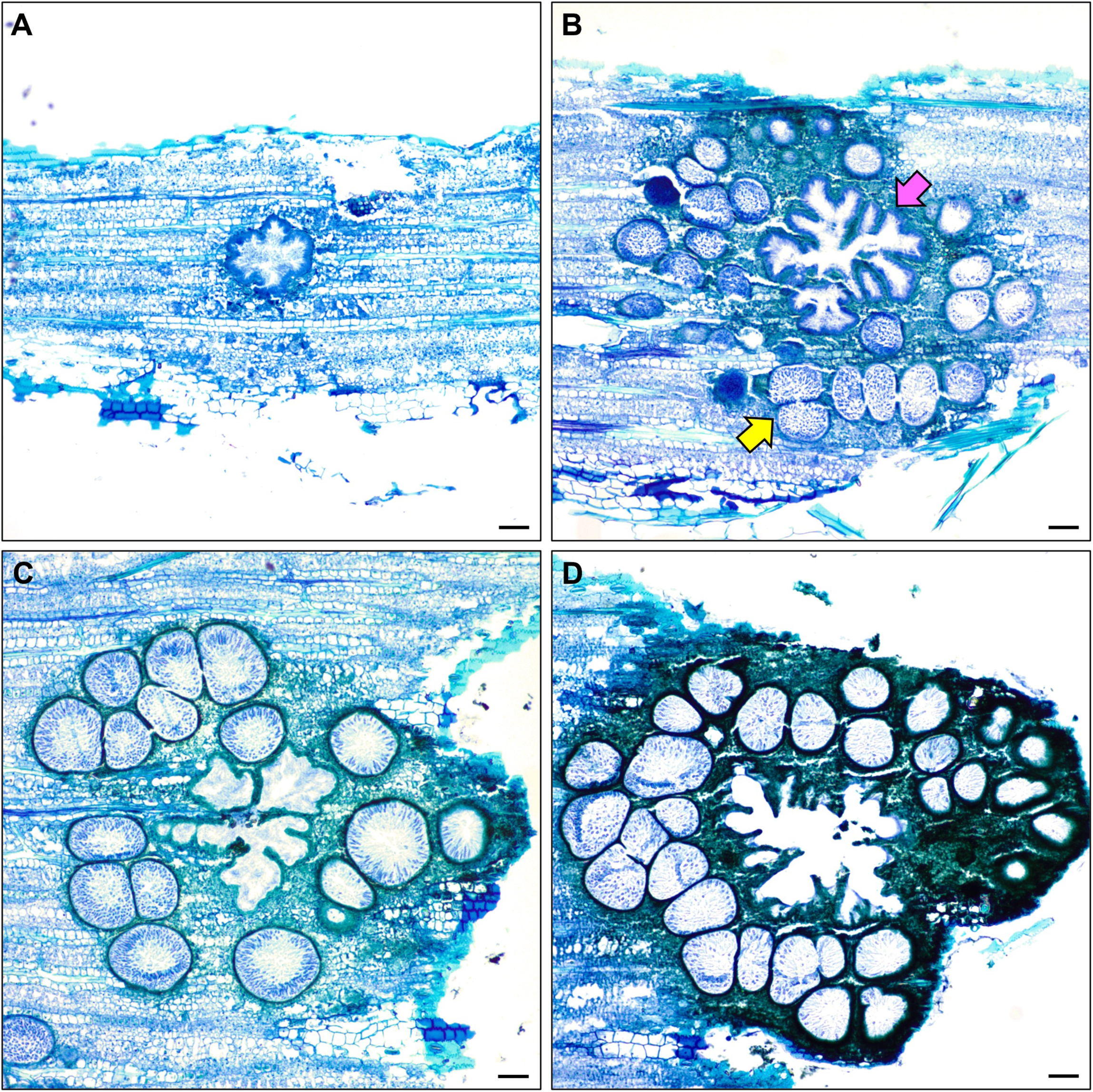
Sections of maize leaf tissue infected with *P. maydis* (lateral orientation). (A) The early stage of stromata development is characterized by a central multilobed pycnidium surrounded by developing ascomata (perithecia). (B-D) As *P. maydis* infection progresses, lobing of the central pycnidium locule (pink arrow) increases and the perithecia (yellow arrow) develop into mature sexual structures. Scale bar represents 10 µm.

### Fungal colonization of maize epidermal cells leads to clypeus development

Upon *P. maydi*s infection, fungal hyphae penetrate and extensively colonize the host epidermal cells. Maize epidermal cells derived from non-infected leaf tissue are translucent from the lateral orientation (Fig. 3A). However, intracellular spread of the hyphae leads to the maize epidermis becoming melanized, resulting in the formation and development of the fungal clypeus (Fig. 3B). The clypeus primarily functions as a shield-like structure for stromatic growth positioned above the vegetative hyphae and the reproductive structures within the developing stromata (Silva-Hanlin and Halin 1998). As the fungal hyphae colonization progresses, the epidermal cells gradually become darker, resulting in the characteristic tar spot lesion formation on the leaf surface (Fig. 3B).

**Fig. 3.**
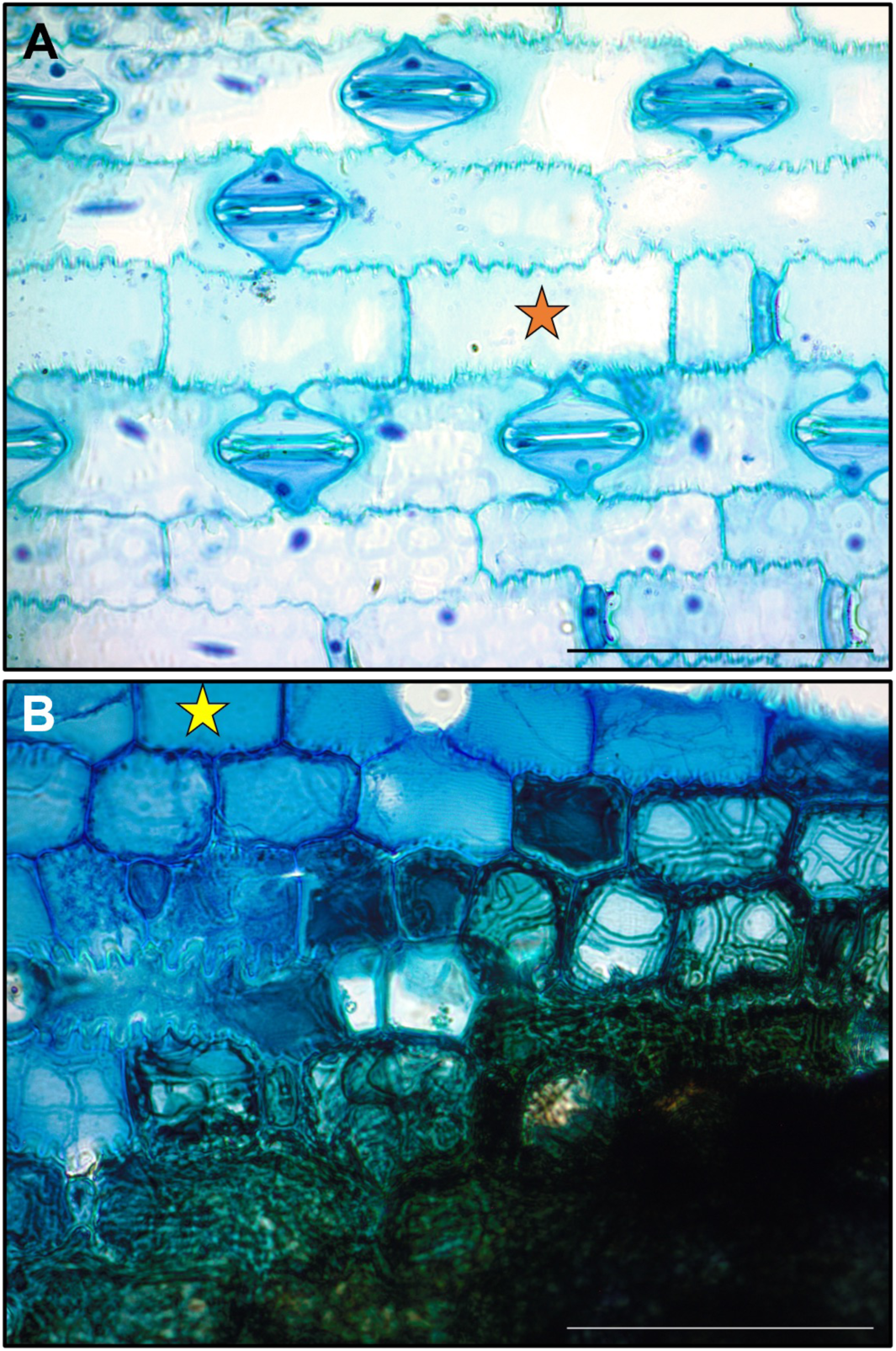
Progression of *P. maydis* disease spread in the maize leaf epidermis. (A) Asymptomatic maize leaf epidermal surface depicting non-infected leaf pavement cells. Stomates are open and free of debris. (B) In a symptomatic leaf, a mix of non-infected and infected epidermal cells with varying concentrations of vegetative hyphae are observed. As cells fill with hyphae, they become obscured and melanized and the mature clypeus forms. Orange star shows a healthy epidermal cell; yellow star shows an uncolonized epidermal cell in an infected leaf. Scale bar represents 10 µm.

In the longitudinal orientation, hyphae continue their colonization of the epidermis and occupy most of the host cell. Ultimately, the hyphae amalgamate to form a densely compacted mass of interwoven hyphae that fully occupies the epidermal cell (Fig. 4, blue arrows). The epidermal wall appears to remain intact until the later stages of colonization.

**Fig. 4.**
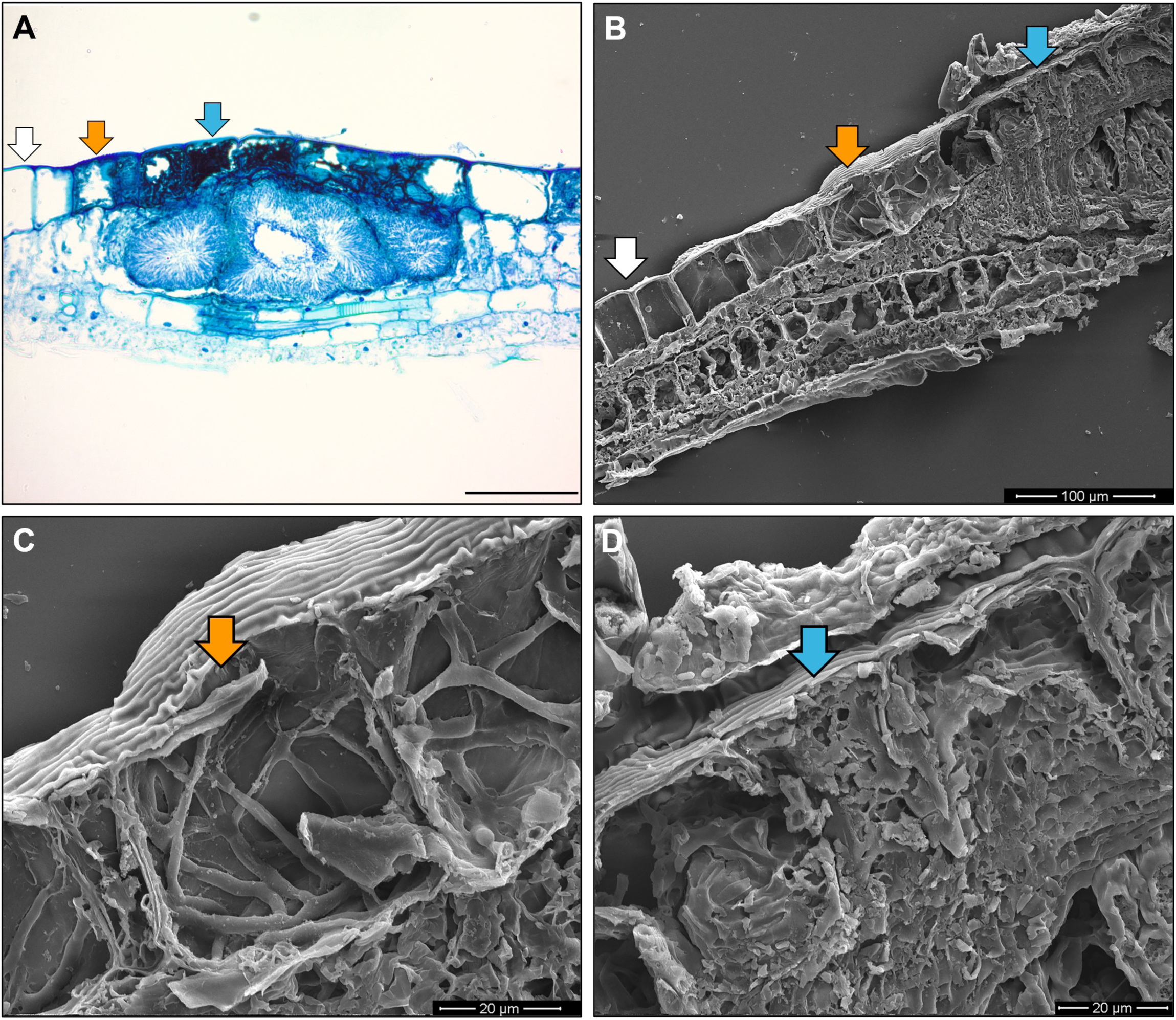
*P. maydis* clypeus formation within maize epidermal cells viewed in the longitudinal orientation. (A) *P. maydis-*infected maize leaf stained with Toluidine Blue. (B) Scanning Electron Microscopy (SEM) image of a *P. maydis-*infected maize leaf. (C) Epidermal cells containing branched, septate hyphae of *P. maydis*. (D) A mass of hyphae develops and occupies the internal epidermal cell. For panels A to D, white arrows indicate non-colonized epidermal cells, orange arrows indicate epidermal cells that are partially colonized by fungal hyphae, and blue arrows indicate epidermal cells that are fully colonized by hyphae forming a dense matted body. Scale bar in A represents 10 µm.

It is worth noting that the development of the clypeus appears not to extend beyond the boundaries of the reproductive tissues (Fig. 4A-B). Conversely, the epidermal cells outside the clypeus do not exhibit the presence of hyphae (Fig. 4A-B, white arrow).

### Fungal hyphae of P. maydis progress through the leaf and develop into pycnidia independent of the host vasculature

As the *P. maydis* clypeus continues to develop within the leaf epidermis, a specialized structure that produces asexual spores known as the pycnidium forms within the mesophyll, proximal to the original site of infection. During the initial stages of fungal infection, the pycnidium initially develops into a flask-shaped structure that progresses into a clover-like structure (Fig. 5A). As the disease progress and the pycnidium matures, it transitions into a multilobed locule (Fig. 5A-B).

**Fig. 5.**
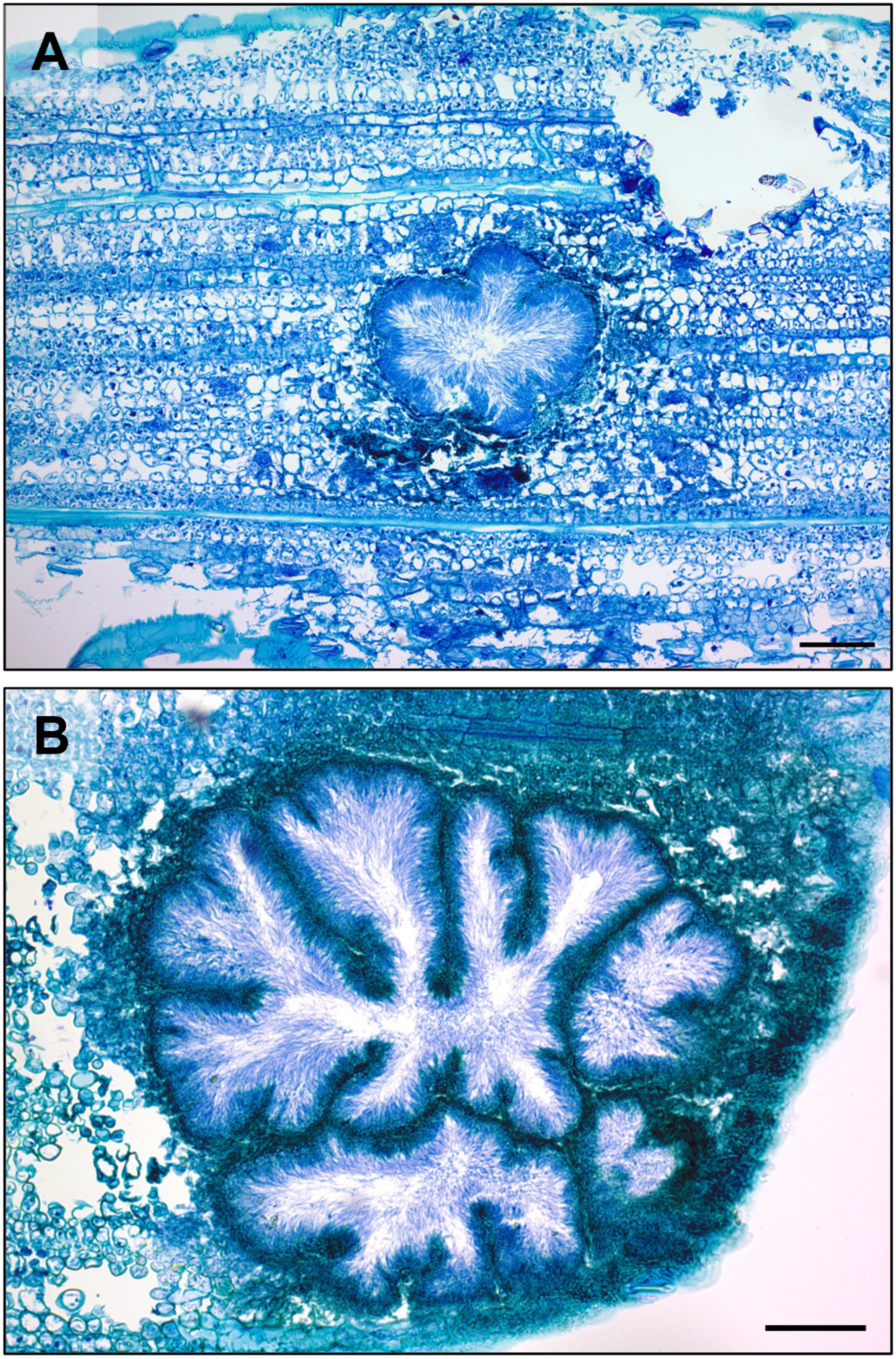
Lateral view of the *P. maydis* pycnidia in the leaf mesophyll. (A) Infected tissue with a clover-shaped central pycnidium in the early stages of *P. maydis* development. (B) The central pycnidium matures into a multilobed pycnidium. Scale bars represent 10 µm.

In the longitudinal orientation, the pycnidium develops from one side of the mesophyll (Fig. 6A). Initially, only a spherical locule is visible. As the pycnidium develops, it expands further into the mesophyll, developing around the host vasculature, transitioning from a spherical shape to a multilobed structure (Fig. 6B). Near the point of infection, a single apical ostiole emerges, through which the asexual spores release (Fig. 6C). As the pycnidium matures, it gradually penetrates deeper into the leaf tissue, while a secondary clypeus develops on the opposite epidermis (Fig. 6D). Though hyphae frequently invade the bundle sheath cells, the vascular tissue, including the xylem and phloem, remain largely uncolonized (Fig. 6B). As the infection progresses and asexual spores are released, empty pycnidia chambers remain in the leaf tissue. The pycnidium undergoes further development along the longitudinal axis rather than the transverse axis, corresponding with the longitudinal expansion of the tar spot lesion on the leaf surface (Fig. 6D).

**Fig. 6.**
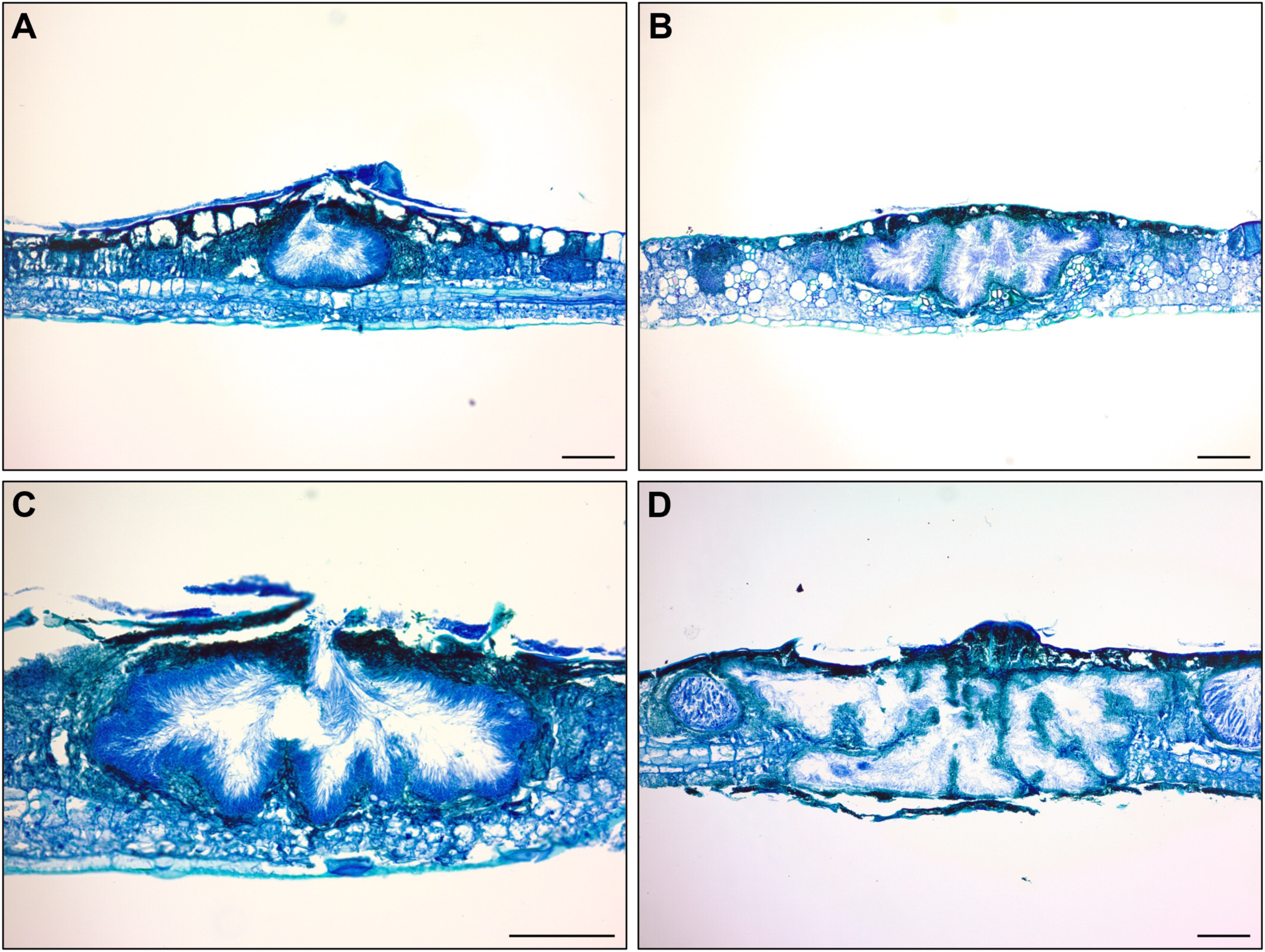
Longitudinal view of *P. maydis* pycnidium within a stromata. (A) The central pycnidium develops beneath the clypeus, often penetrating halfway through the leaf tissue. (B) The pycnidium expands and becomes multilobed, (C) penetrates the abaxial surface of the maize leaf, and (D) develops into both the abaxial and adaxial sides of the leaf. Scale bars represent 10 µm.

After being expelled through the ostiole, asexual spores are present on the leaf surface (Fig. 7A). These asexual spores measure between 7-10 µm and are often filiform in shape (Fig. 7B-D). The spores form a dome-shaped structure located above the stroma (Fig. 7E-F).

**Fig. 7.**
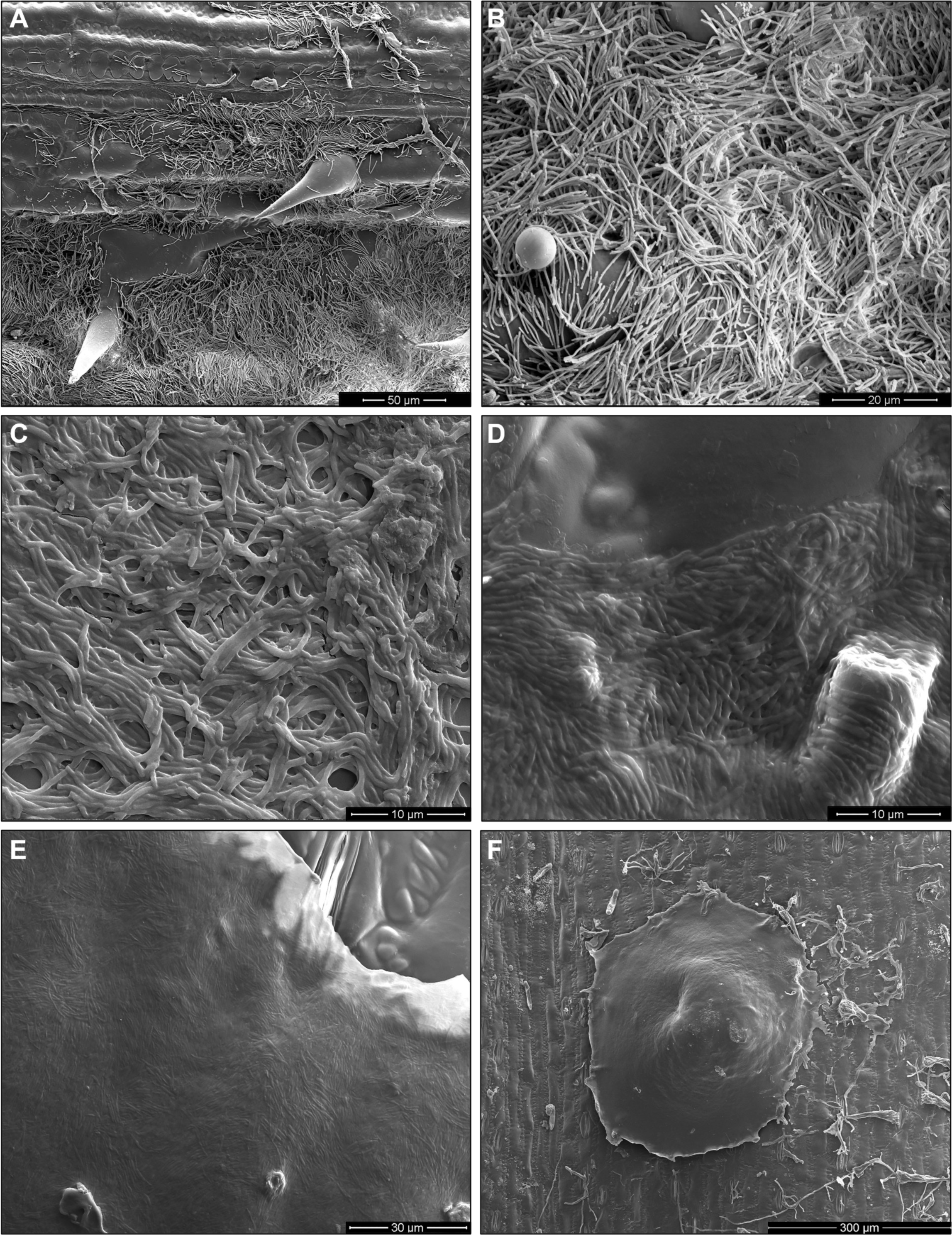
Asexual spores of *P. maydis*. (A) and (B) Asexual spores on maize leaf surface. (C) Mass of asexual spores. (D) and (E) Asexual spores forming a sheet on the surface of epidermal cells. (F) A young stromata with asexual spores exuding from the pycnidium beneath.

Fungal hyphae move through the leaf and emerge from stomata on the leaf surface (Fig. 8). Finger-like projections emerge from the stomata (Fig. 8A-D). Inside the leaf tissue, vegetative hyphae move through the mesophyll toward adjacent stomatal chambers (Fig. 9A-C). Hyphae are observed within the stomatal chamber (Fig. 9 B-E).

**Fig. 8.**
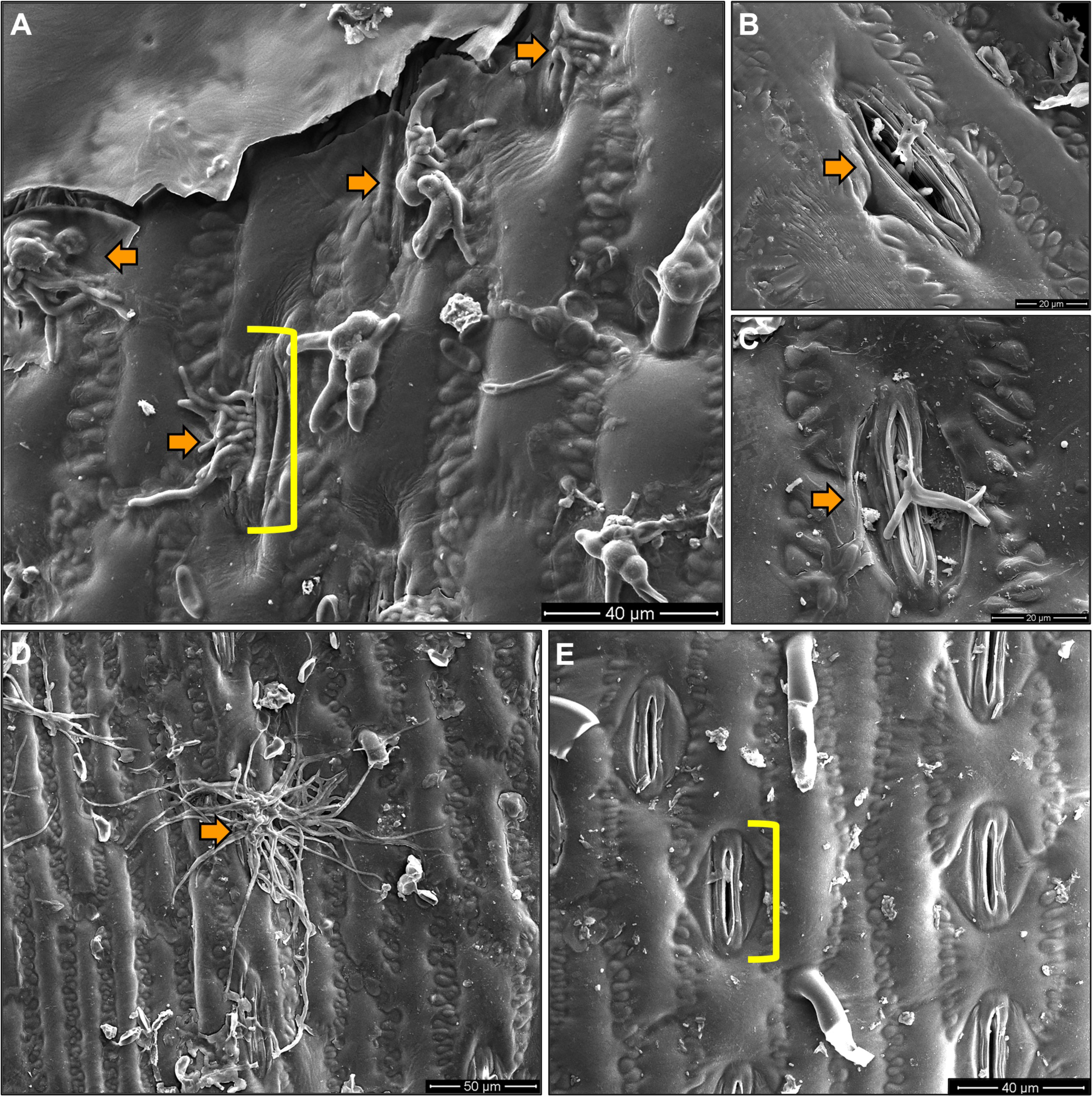
*P. maydis* hyphae emerge from stomates on the maize leaf surface. (A-D). SEM micrographs of *P. maydis* hyphae emerging from stomates on the maize leaf surface. Orange arrows label the hyphae; yellow brackets label the stomate. (E) Healthy leaf tissue with stomate.

**Fig. 9.**
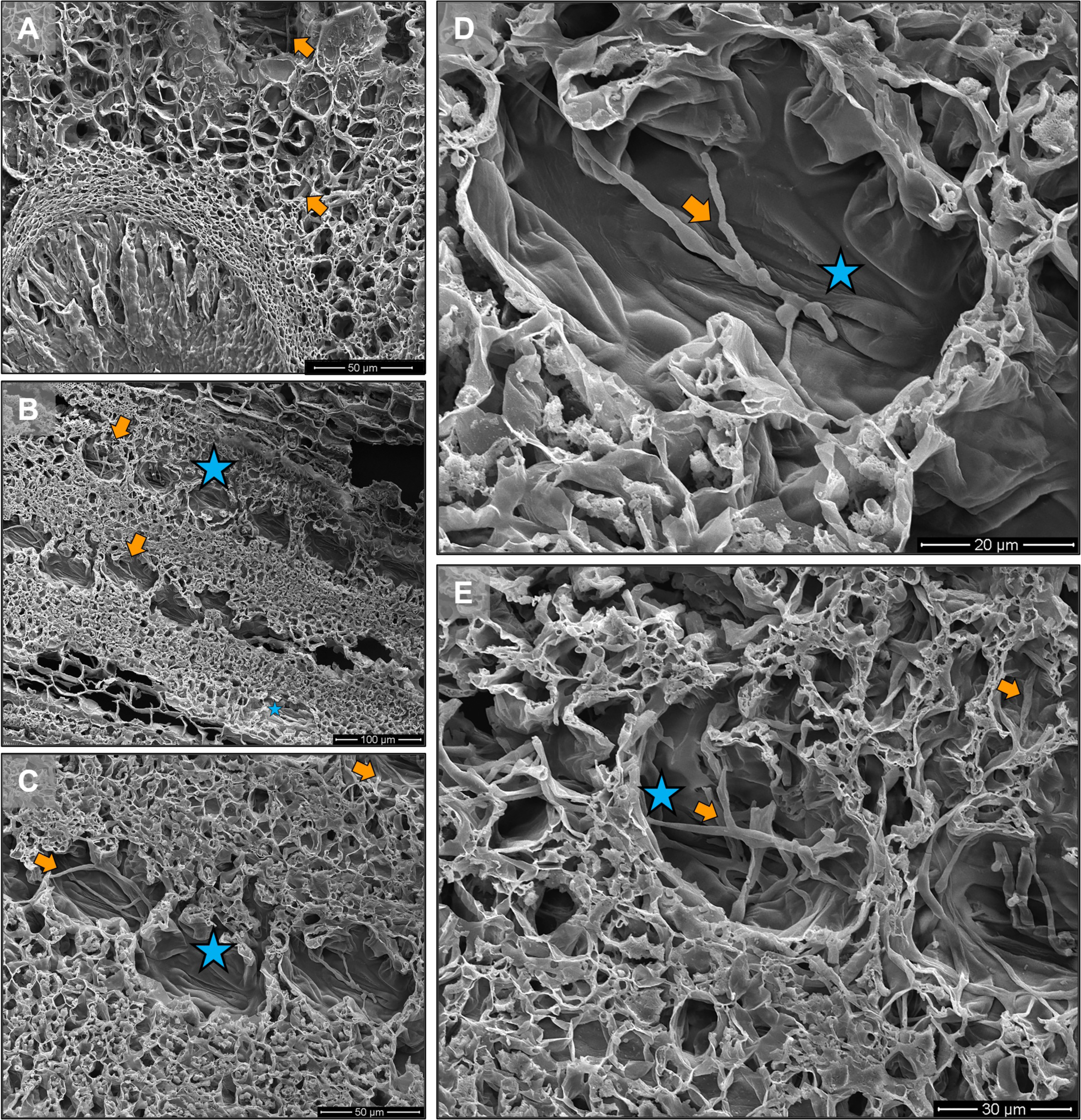
Depiction of the epidermis (abaxial) of *P. maydis*-infected maize leaf tissues. (A) *P. maydis* vegetative hyphae within the mesophyll. (B-E) Hyphae colonize stomatal chambers toward stomates. Orange arrows label the hyphae; blue stars label the stomatal chambers. For panel B, stomatal chambers are observed as a line of wells in the leaf tissue.

### Sexual fruiting body development

The guard cells in non-infected tissue remain visible and unobstructed (Fig. 10A). As the colonization of stomata progresses, hyphae protrude between the guard cells (Fig. 10B). Additional hyphae remain within the leaf tissue, forming a lining along the walls of the stomatal chamber, likely signifying the initiation of perithecium development (Fig. 10C). The hyphae both inside and outside the stomate increase in density (Fig. 10C). Above the stomate, a cluster of hyphae form, while hyphae connect the interior and exterior of the leaf between the two guard cells (Fig. 10C).

**Fig. 10.**
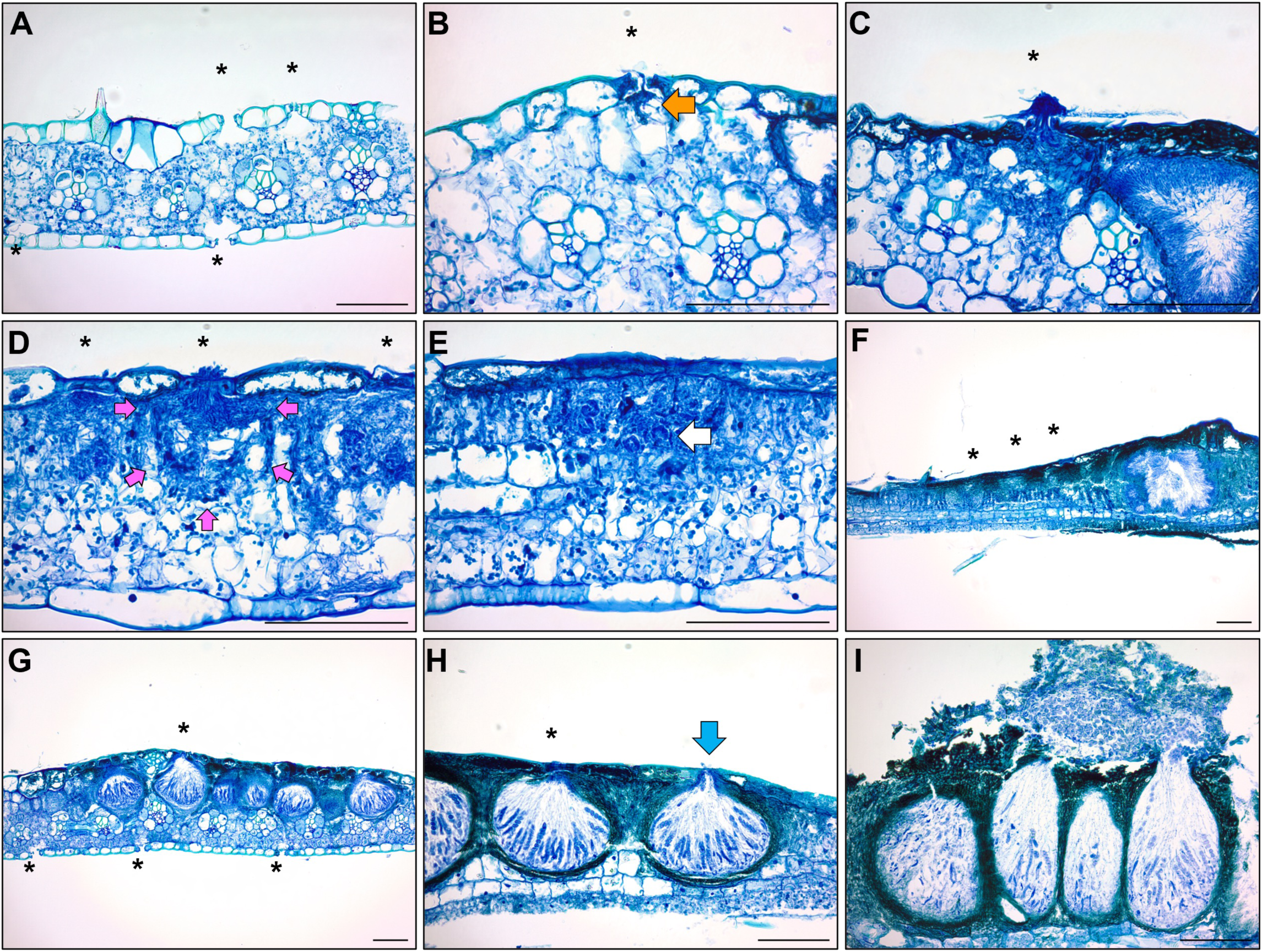
Sexual fruiting body development of *P. maydis* on a maize leaf. (A) Healthy leaf tissue, (B) Early stages of hyphae (orange arrow) entering stomates on the adaxial side of the leaf. The adaxial epidermis is filling with hyphae, promoting clypeus development. (C) A large mass of hyphal material is on the outside of a pair of guard cells as a mass grows within the chamber below. (D) Hyphae line the stomatal chamber and the perithecium develops. Pink arrows depict the outline of the developing perithecium. (E) Hyphae differentiate into crozier hooks (white arrow) initiating the base of the asci. (F) Perithecium are present in the stomatal chambers directly below the stomates adjacent to the central pycnidium. (G) Asci formation is initiated. (H) Ostiole forms in the clypeus (blue arrow) upon maturity of the perithecium (I), and the spores release through the ostiole. Black stars represent stomates.

The perithecium walls become apparent at the periphery of the stomatal chamber (Fig. 10D). The hyphae continue to colonize the perithecia and eventually form croziers (Fig. 10E). During this stage, the hyphae cease to exit the stomate, and the perithecium wall develops around the maturing asci and ascospores. The adjacent stomata continue to develop, and the perithecia walls mature progressively from the base to the top, growing into the mesophyll while circumventing the vascular bundles (Fig. 10F). In the intermediate stage of perithecium development, perithecia reside adjacent to each other, mirroring the pattern observed in panel F (Fig. 10G). Before the release of ascospores, the walls between the perithecia become interwoven with hyphae (Fig. 10H). Within the surface of the clypeus, the ostiole begins to take shape (Fig. 10H) and is the exit point for ascospore release (Fig. 10I).

The perithecia, similar to pycnidia, form around the vasculature, which is not colonized by *P. maydis* hyphae (Fig. 11A-B). Within the leaf mesophyll of the stroma, there can be two perithecia, with one positioned towards the upper surface of the leaf and the other towards the lower surface. Both perithecia are enclosed by the clypeus and possess an ostiole. The development and orientation of the perithecia are directed towards the opposing leaf epidermis (Fig. 11A-B).

**Fig. 11.**
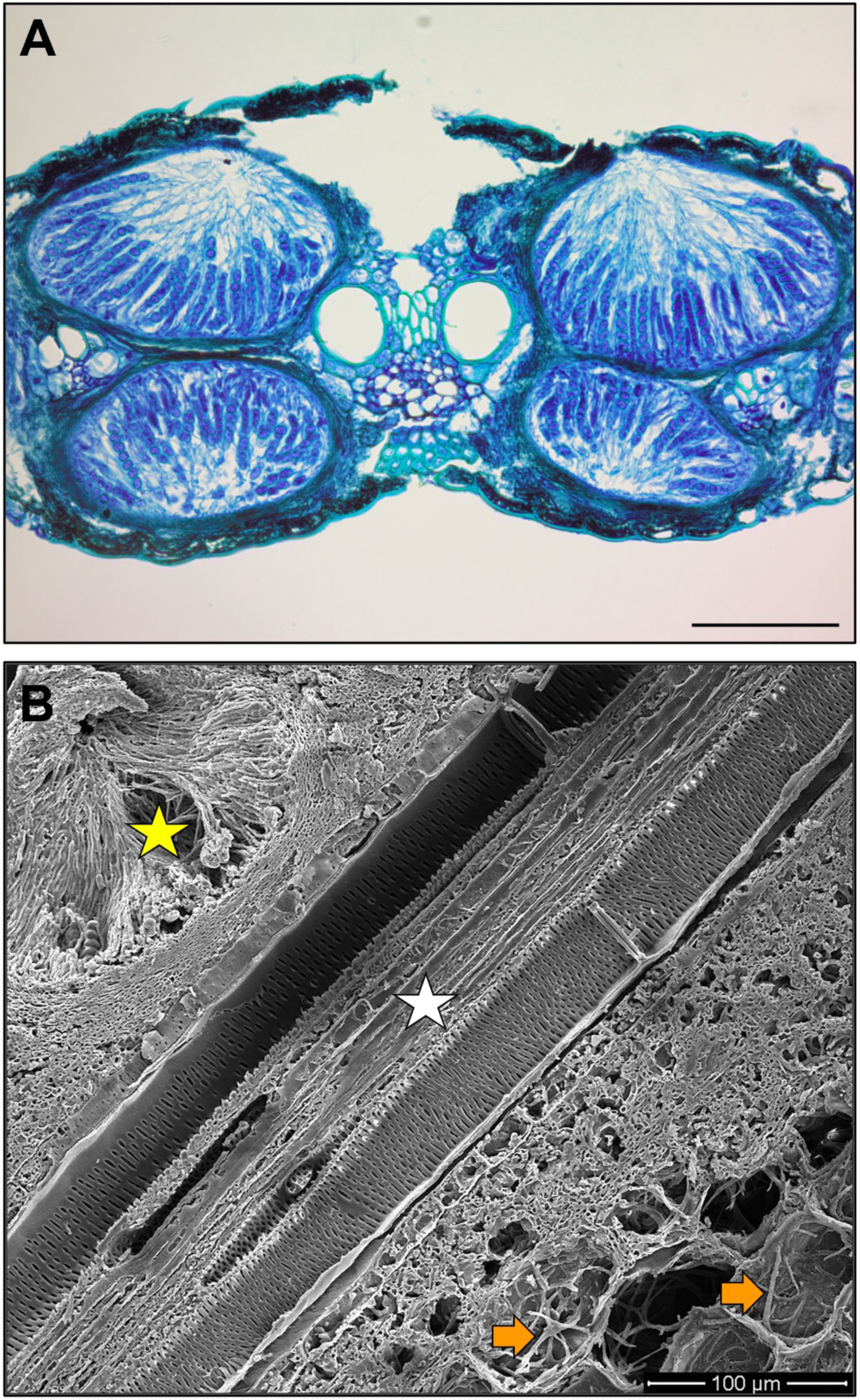
*P. maydis* perithecia form around the vasculature but the fungus does not appear to colonize xylem or phloem. (A) *P. maydis* colonizes the mesophyll space between the upper and lower epidermis but major and minor veins are not colonized. (B) In a longitudinal SEM image, the upper leaf surface has a well-developed perithecium (yellow star) and hyphae colonize the lower leaf surface (orange arrow), but the vasculature (white star) remains free of *P. maydis*. Light micrograph scale bar represents 10 µm.

### Ascospore release is through the ostiole and not through the stomate

As the perithecium matures, the chamber walls gradually close (Fig. 12A). When the perithecia reach maturity they swell, forcing the ascospores through the ostiole (Fig. 12B-C). Once released, the ascospores likely germinate and develop appressorium on the leaf surface, apparently separate from the maternal stroma (Fig. 12D).

**Fig. 12.**
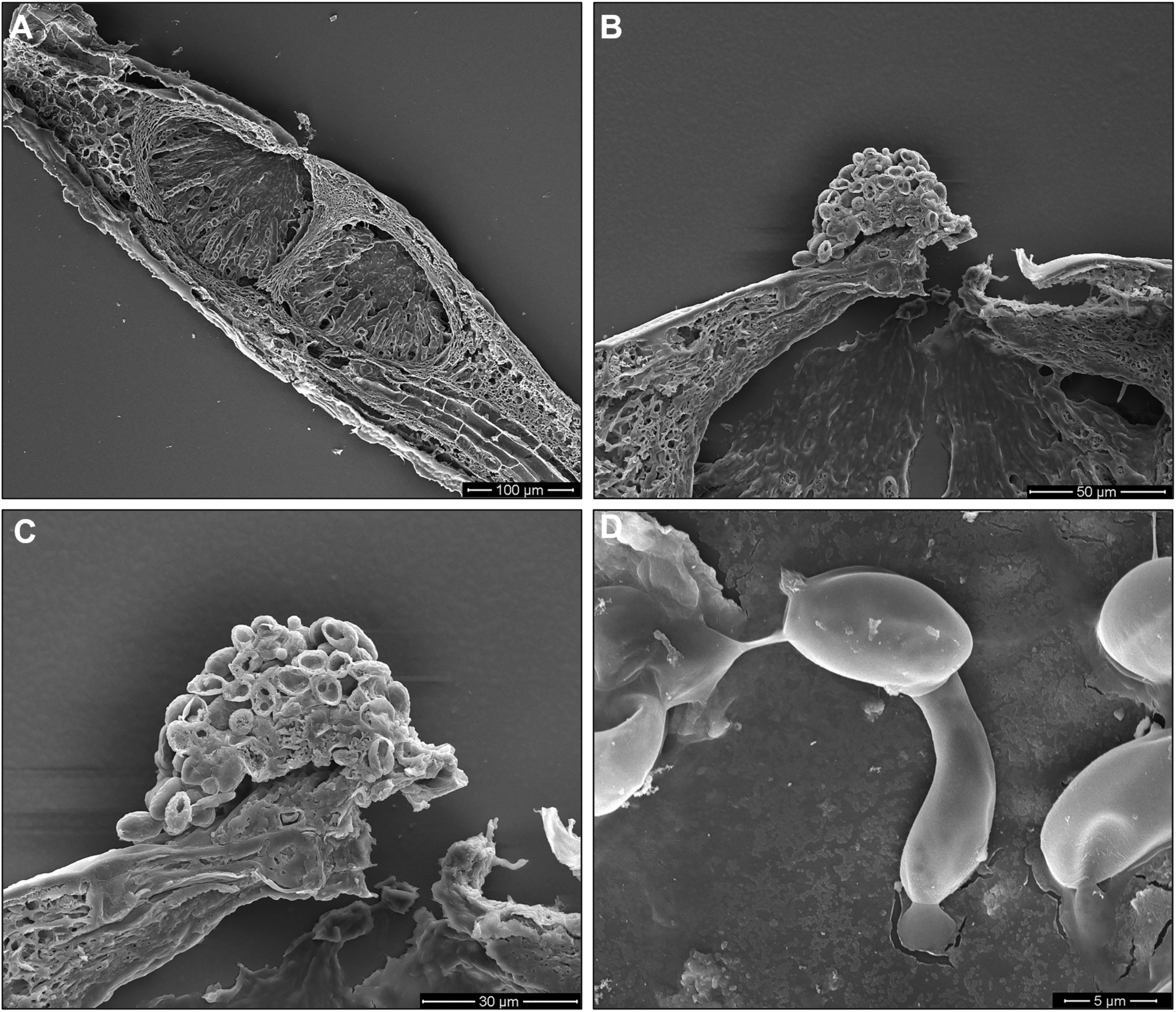
SEM micrographs of the sexual reproductive structures of *P. maydis* in an infected maize leaf. (A) Longitudinal 20 µm section of two perithecia under a closed clypeus. (B and C) Ascospores are emerging from the ostiole. (C) Shows a higher magnification image of B. (D) Spore germinating on the maize leaf surface.

### Hyperparasites of Phyllachora maydis: Fish eye

A characteristic fish eye lesion is occasionally observed as part of tar spot disease progression (Hock et al. 1992, 1995; McCoy et al. 2019; Rocco da Silva et al. 2021; Telenko et al. 2020; Valle-Torres et al. 2020). Of the four hybrid corn fields we sampled, fish eye lesions were present on leaves from three fields. Lateral and longitudinal views revealed that *P. maydis* disease structures were present in these samples (Fig. 13A-D).

**Fig. 13.**
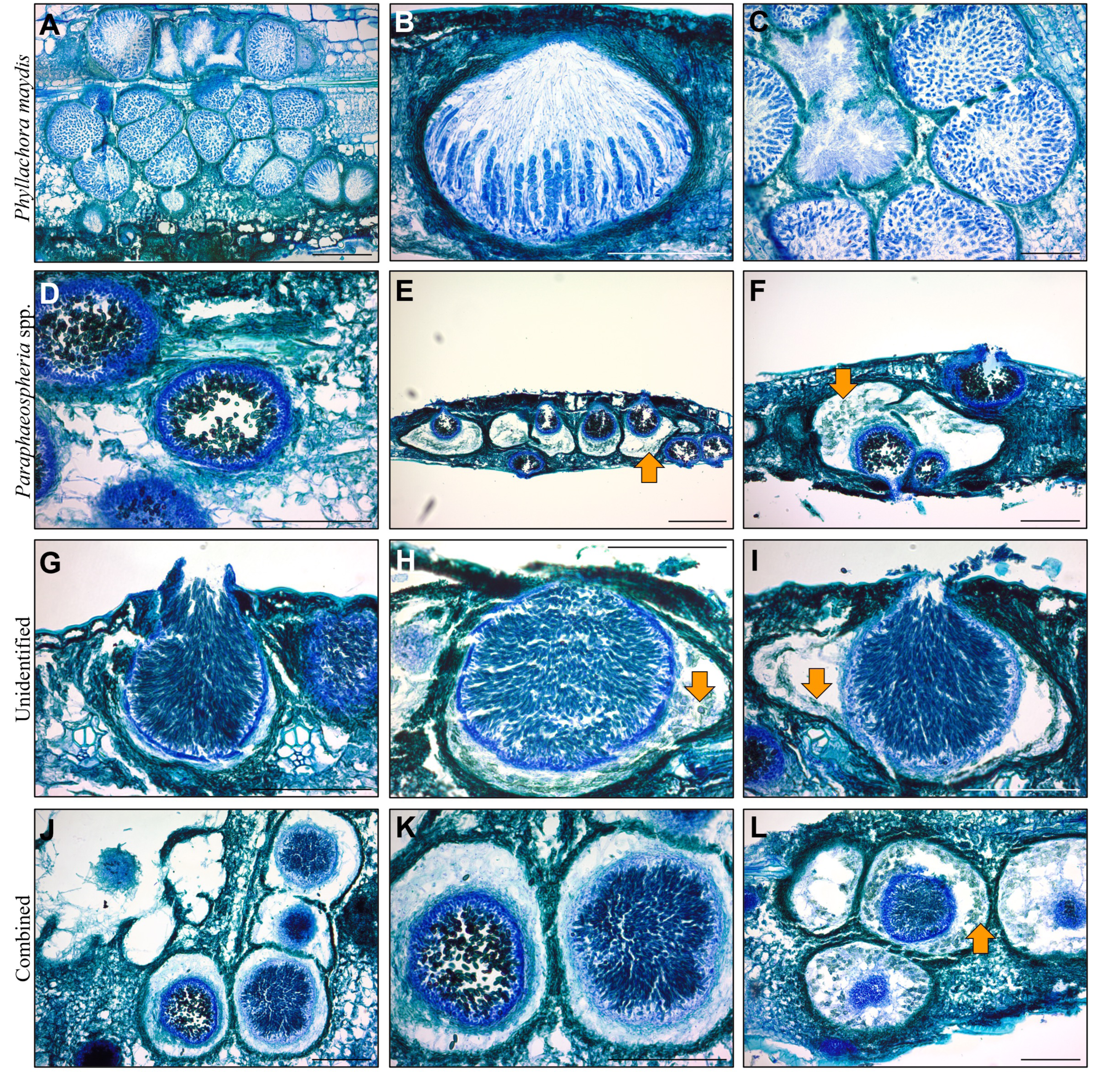
Different spore types found in fish eye lesions. (A-C) *P. maydis (*D-F) Suspected *Paraphaeospheria* spp. (G-I) Third, unidentified fungus associated with fish eye. (J-L) Show the two parasitic pathogens with fully formed pycnidium inside the *P. maydis* perithecium. The orange arrows point to parasitized *P. maydis* ascospores. Scale bar represents 10 µm.

In addition to *P. maydis*, two different fungal spores and structures were also present in fish eye samples. These additional spores existed within the *P. maydis* perithecium (Fig. 13E-I, Supplementary Fig. 5-6). Both of these hyperparasites exploit the perithecium and ostiole of *P. maydis* to develop and subsequently release their own spores.

To identify the additional spores in the fish eye samples, we sequenced genomic DNA extracted from tar spot stroma exhibiting *P. maydis* only (samples 1, 2, 3, 7, 8, 9) and stroma with the presumed *Paraphaeosphaeria* spp. and the unknown species together (4, 5, 6, 10, 11, 12) (Fig. 14). Samples 2, 5, and 9 failed during Illumina sequencing and could not undergo fungal microbiome characterization. Of the Illumina sequences obtained from each remaining sample 3-11% mapped to the B73 *Zea mays* genome and 15-72% mapped to the PM02 *P.maydis* genome. The remaining reads were used for DIAMOND and MEGAN BLAST analysis. The mycoparasitic genera *Paraphaeosphaeria* and *Coniothyrium* were present in nearly all samples (Fig. 14). *Paraphaeosphaeria* spp. were highly abundant in samples 1, 3, 4, and 6; absent in samples 7 and 8; and lowly abundant in samples 10, 11, and 12 (Fig. 14). *Coniothyrium* spp. sequences were identified in all samples except sample 7. It is unclear if these *Coniothyrium* spp. are still taxonomically classified as such, or if these species are now within the genus *Paraphaeosphaeria* (Verkley et al. 2014). *Stagonospora* spp. were found in all samples sequenced. In a previous microbiome study, *Neottiosporina paspali,* a Stagonospora-like fungus, was identified as being significantly more abundant in fish eye lesions than that found in tar spot only symptomologies. *Monographella* spp. were not identified in any of the samples, however, *Fusarium* spp. were identified in all samples. *M. maydis* (syn*=Michrodochium maydis*), was once believed to cause fish eye symptoms, but isolates from Mexico were recently molecularly identified as *Fusarium* spp. within the *F. equiseti-incarnatum* and *F. sambucinum* species complexes (Luis et al. 2023).

**Fig. 14.**
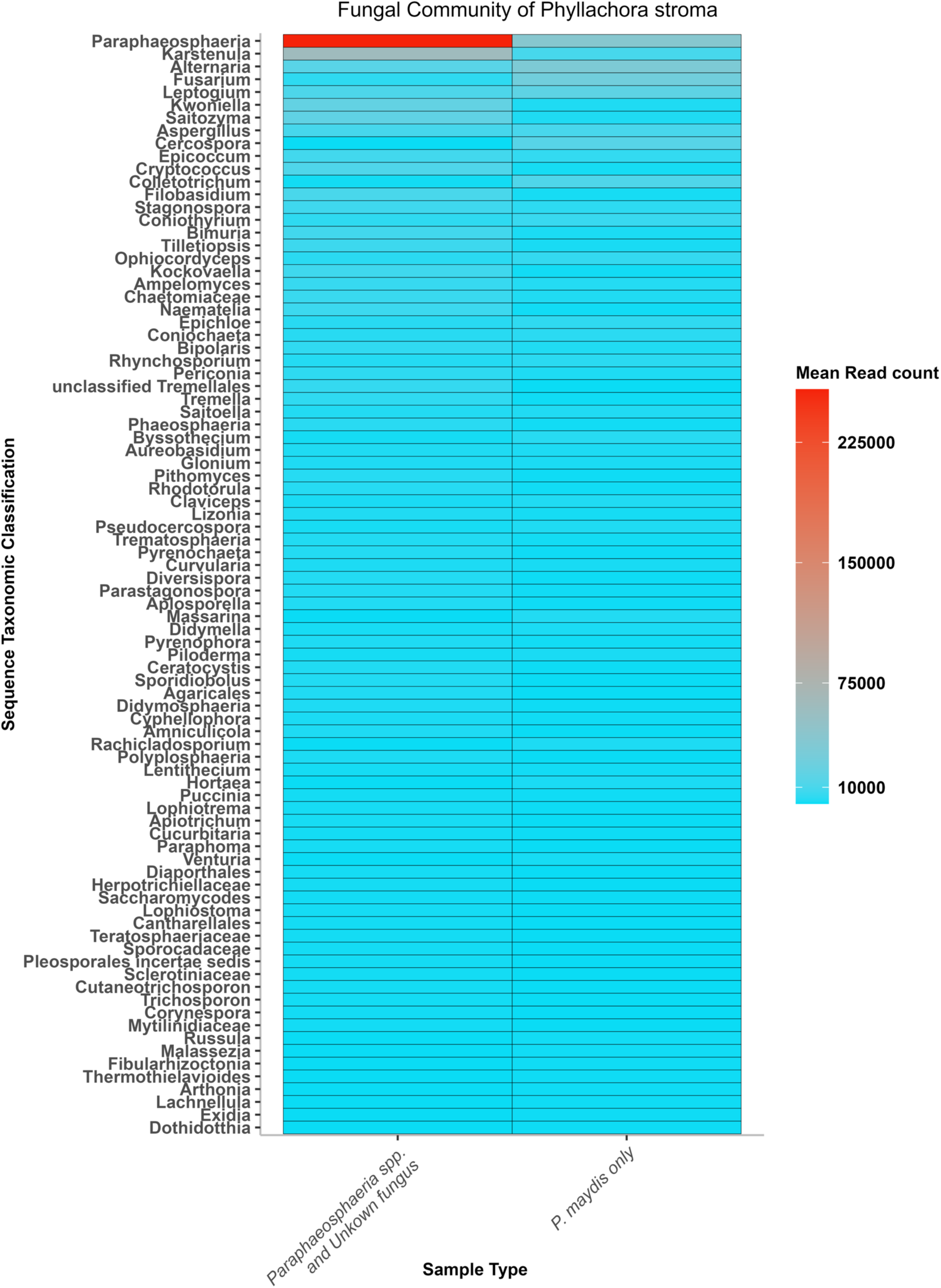
Fungal community of *P. maydis* stroma. Leaves sampled are denoted on the x-axis with fungal Genera identified from Illumina shotgun sequencing via DIAMOND and MEGAN analysis on the y-axis for each sample. Only genera with over 1,000 reads in a given sample were visualized. Shading of tiles indicates abundance of each genera within each sample from low abundance (turquoise blue) to high abundance (red).

## DISCUSSION

Tar spot disease has caused significant yield loss in Central and South America for many decades (Hock et al. 1992, 1995), and since its emergence in the U.S. in 2015, has continued to limit production throughout corn-producing regions of the country. While work characterizing *P. maydis* infection has not been reported, earlier studies suggested that other *Phyllachora* spp. formed perithecia and pycnidia (Parbery 1963b, 1967, 1963a; Orton 1944) but also colonized vascular cells (Gabel 1989). How *P. maydis* spread within the leaf, emerged from the leaf, and developed new stromata on the leaf was not clear. Additionally, components of the fish eye complex in the U.S. were not well described. Here we show how *P. maydis* infects maize leaves, reproduces, and disseminates within the stroma, the leaf surface, and amid maize plants. We observed a consistent pattern of infection in different samples collected across fields. We were unable to observe initial penetration of the fungus due to the selection of mature stroma.

However, similar to previous work with other *Phyllachora* spp. (Parbery 1963b, 1963a, 1967; Orton 1944), we found that each stromata contained a central pycnidium that produces asexual spores, perithecia that surround the central pycnidium, and fungal hyphae that colonize epidermal cells. Epidermal cells colonized with hyphae become melanized, forming the clypeus. Asexual spores are released through an ostiole of the pycnidia, and a cap of spores forms over the stroma. Hyphae emerge through stomates and perithecia with sexual spores develop in the underlying stomatal chambers. Sexual spores are released through an ostiole of the perithecia, promoting reinfection on the same leaf. We hypothesize that multiple perithecia in different stages of development may facilitate the continuous production of *P. maydis* at varying rates, enabling reinfection and resulting in the characteristic elongated tar spot lesion observed along the maize leaf (Supplementary Fig. 7-10G).

Our data suggest disease spread within a leaf is likely due to the release and subsequent germination of ascospores on the leaf surface, rather than the spread of hyphae throughout the leaf. In contrast to a previous report characterizing *Phyllachora* spp. on warm season grasses (Gabel 1989), *P. maydis* does not appear to be a vascular pathogen, as we examined 580 stroma samples and the presence of *P. maydis* was absent within the vasculature in all instances. Though fungal hyphae were often observed within bundle sheath tissues, we did not observe fungal hyphae penetration into the vasculature. The breakdown of bundle sheath cells can disrupt photosynthesis, which could lead to the leaf chlorosis commonly observed during severe tar spot disease outbreaks.

### Differing rates of ascospore development may promote disease spread across the leaf

Our results suggest a correlation between perithecium development and the arrangement of stomates in maize. During maize leaf development, parallel veins are formed, interspersed with files of cells containing stomates. The precise spacing between stomates is achieved through asymmetric cellular divisions, ensuring each stomate is positioned exactly one cell apart from neighboring cells (Nunes et al. 2020). Perithecia development followed stomate positions longitudinally along the leaf (Fig. 10, Supplementary Fig. 2-3) and hyphae transitioned from the stroma structure to adjacent empty stomatal chambers (Fig. 9). The chambers near the originating stroma were the first to be invaded, suggesting that ascospores mature at different times throughout the season. Genomic evidence suggests that *P. maydis* is heterothallic, requiring two mating types for sexual reproduction and the formation of ascospores (MacCready et al. 2023). Variable temporal formation of perithecia and maturation of ascospores within the stroma on a single leaf surface may be indicative of separate spatial sexual recombination events. We hypothesize that aerially dispersed ascospores from previous years’ infested material would initiate infection and pycnidium development on the leaf. Spermatial conidia oozed from the pycnidia would provide the complementary mating type for sexual reproduction in receptive pycnidia on the leaf and lead to the formation of the next generation of ascospores. Individual perithecia would then release spores independently from the stroma at various times, promoting a continuous supply of inoculum and ensuring the presence of ascospores throughout the season.

### Stromata are distinct from one another

Upon release, ascospores exhibit germination on non-infected areas of the leaf. Stromata were distinct and not interconnected. This could be due to several factors, such as the field collection method, spores, or the potential inhibitory effects of stromata. In other ascomycetes like *Penicillium paneum*, high concentrations of asexual spores hinder mycelial growth and spore germination (Chitarra et al. 2004). Similarly, *Aspergillus nidulans* produce spores that generate inhibitory substances and reduce local oxygen upon reaching a specific density, affecting conidial germination (Trinci and Whittaker 1968). It is possible that significant levels of asexual spores inhibit germination of other pathogens within the vicinity of the stroma or hinder the early germination of *P. maydis* ascospores within the perithecium or the local stromatic area.

### Does the clypeus limit the size of the stromata?

Previous work investigating several species of *Phyllachora* revealed that the clypeus begins as a separate structure but fuses with the pycnidium and perithecium as the reproductive structures develop, forming a single entity (Parbery and Langdon 1963). Observations on the growth of *P. graminis* suggested a correlation between clypeus development and growth of the perithecia and pycnidium as the clypeus does not appear to extend beyond the boundaries of the perithecia (Parbery 1963b). Our results are consistent with this and also show that in *P. maydis,* perithecia envelop the central pycnidia, causing these structures to conjoin and form the stromata. Together these data suggest that clypeus development is closely linked to the formation and growth of the perithecia and pycnidium.

### Fish eye lesions are likely composed of three different pathogens

Fish eye lesions are often but not always present in tar spot infected fields. When present, such lesions are commonly known as the tar spot complex. In Latin America two fungi are present in fish eye lesions – *P. maydis* and *M. maydis*. Previous work suggested that *M. maydis* is not a component of fish eye lesions in the U.S. (McCoy et al. 2019). Consistent with this, we did not observe *M. maydis* associated with tar spot lesions in our samples. Instead, our sequencing and morphological identification suggested that *Paraphaeospheria* spp. is a hyperparasite of *P. maydis* associated with fish eye, and a third unknown pathogen is also present in Indiana samples. Our finding of this previously undescribed third fungus underscores the complexity of tar spot and fish eye lesions and shows how much more we have to learn about tar spot.

### Conclusions

Our study highlights significant cellular aspects of *P. maydis* infection and sheds light on the disease cycle and fish eye lesions. Our findings that *P. maydis*, but not the hyperparasites, is the primary fungus present in tar spot but that the hyperparasites are associated with fish eye lesions suggests that tar spot disease and tar spot disease complex are two separate diseases. Our work can inform the development of new approaches to managing tar spot, which could help mitigate the significant economic losses caused by this disease. In addition, this knowledge leads to new questions and enables future studies of *P. maydis* biology and the molecular mechanisms of disease progression. For example, questions regarding the role of leaf development in tar spot infection and the relationship between the clypeus and underlying reproductive structures remain to be addressed. Given that hyphae emerge from stomates, could stomatal resistance strategies be effective against *P. maydis*? How does *P. maydis* develop seemingly unnoticed within susceptible maize leaves? We hypothesize that *P. maydis* uses virulence proteins (effectors) to modulate host immune responses, thereby facilitating the infection process (Helm et al. 2022; MacCready et al. 2023; Rogers et al., 2023). Our work provides the foundation for further research to uncover the molecular mechanisms underlying *P. maydis* cellular invasion and colonization.

## ACKNOWLEDGEMENTS

This work was funded by the Purdue AgSeed program to AIP and DT, an Indiana Corn Marketing Council Graduate Assistantship to DC, and partial support from the Corn Marketing Program of Michigan and Project GREEEN-Michigan’s plant agriculture initiative. This research was also funded, in part, by the United States Department of Agriculture, Agricultural Research Service (USDA-ARS) research project 5020-21220-014-00D. The funding bodies had no role in designing the experiments, collecting the data, or writing the manuscript. All opinions expressed in this paper are the author’s and do not necessarily reflect the policies and views of USDA. USDA is an equal opportunity provider and employer. We thank members of the Iyer-Pascuzzi, Helm, Telenko and Chilvers labs for critical reading of the manuscript and Chloe Caldwell for technical help.

## SUPPLEMENTARY INFORMATION

**Supplementary Fig. 1: Different orientation planes show the direction of cuts across the leaf blade.** (A) Transverse, or cutting the longitudinal axis at 90-degree angle. (B) Longitudinal, cutting along the long axis of the leaf blade perpendicular to the transverse plane. (C) Lateral plane, cuts the leaf horizontally top to the bottom, and (D) three planar cuts that together make a 3-demensional figure from 2-deminsional figures.

**Supplementary Fig. 2: Transverse (A-K) and longitudinal (L-V) serial sections of stromata early in development.** (A) Leaf with stroma. (B) Section of leaf with stroma showing the direction of serial cuts (arrow). C’ indicates the first cut; K’ indicates the last. (C-K) Serial sections through one stromata. Because this is in the transverse plane, cell lines can be followed. The same cell line in each image (C-K) is indicated by an orange arrow. The blue diamonds are positioned over stomates, and the black arrow points to hyphae within a cell. Image C is on the outer periphery of the stroma, while Image I is in the center of the stromata. A well-developed clypeus is visible and immature pycnidia with conidiospores are developing. The young pycnidium is globose in shape, with a second lobe forming, and occupies the upper surface of the leaf while growing around the veins. (L) Leaf with stroma. (M) Section of leaf with stomata showing the direction of serial cuts (arrow). N’ indicates the first cut; V’ indicates the last. (N-V): Serial sections through one stromata. Hyphae in the upper epidermis form the clypeus and the side wall of the pycnidium emerges in deeper sections. By section Q, the pycnidium is no longer globose but has developed lobes. In the center of the stromata emerges (S-T), a long pycnidium with a clypeus, mature conidiospores, and a central ostiole are present. This long pycnidium remains on the upper side of the leaf and is outside the vasculature.

**Supplementary Fig. 3: Transverse (A-K) and longitudinal (L-V) serial sections of stroma in mid-disease development.** (A) Leaf with stroma. In mid-development, the disease has spread to more of the leaf. The stoma has continued to develop and have increased more in the longitudinal than the transverse direction but do not overlap. (B) Section of leaf with stromata showing the direction of serial cuts (arrow). C’ indicates the first cut; K’ indicates the last. (C-K) Serial sections through one stromata. The blue diamonds are positioned over stomates. Sexual structures are present and an upper and lower clypeus in the epidermis. Moving through sections, perithecia surround the vasculature. The ostiole forms on the leaf abaxial side. (H-J) Closer to the center of the stromata, the pycnidium has developed. (L) Leaf with stroma. (M) Section of leaf with stomata showing the direction of serial cuts (arrow). N’ indicates the first cut; V’ indicates the last. The blue diamonds are positioned over stomates. (N-V): Serial sections through one stromata in the longitudinal orientation. (L-V), additional perithecia develop on the stromata periphery. The central pycnidium continues to develop multiple connecting lobes with a single opening through the adaxial clypeus. In longitudinal sections the increase in stromata length is apparent. This follows what is observed in the transverse sections: the disease travels along the cell lines with minimal transverse spread. Moving through serial sections reveals a developmental series of the perithecium, with the most mature closest to the pycnidium and the less mature at the outer edges of the stroma.

**Supplementary Fig. 4: Transverse (A-K) and longitudinal (L-V) serial sections of stroma in late disease development.** (A) Leaf with stroma. In late development, the disease has spread throughout the leaf. The stroma has continued to develop and have increased more in the longitudinal than the transverse direction but do not overlap. (B) Section of leaf with stromata showing the direction of serial cuts (arrow). The disease follows the cell lines in the monocot leaf. C’ indicates the first cut; K’ indicates the last. (C-K) Serial sections through one stromata. The orange arrow points to the vasculature. Stroma do not form on top of each other, but smaller stroma form closer to larger stromata. An upper and lower clypeus stays directly over the reproductive structures. The vasculature is not occupied by *P. maydis*. The central multi-lobed pycnidium is present from one epidermis to the other (I-J). (L) Leaf with stroma. (M) Section of leaf with stomata showing the direction of serial cuts (arrow). N’ indicates the first cut; V’ indicates the last. (N-V): Serial sections through one stromata in the longitudinal orientation. The stromata has increased in length. In the center (S), a multi-lobed pycnidium is visible around the vasculature system (U). (S-V) Pycnidia are flanked by many perithecia.

**Supplementary Fig. 5: *P. maydis* reproductive structures colonized by parasitic pathogens.** (A) Remnants of *P. maydis* pycnidium and perithecium with parasitic pycnidium inside. (B) *P. maydis* perithecia colonized by parasitic pathogens. The orange arrow indicates asci with ascospores still attached, the green arrow is suspected *Paraphaeosphaeria* spp., the yellow arrow indicates an unidentified parasite, purple arrow shows degrading *P. maydis* spores.

**Supplementary Fig. 6: SEM of different spore types found in fish eye lesions.** The blue arrows point to *P. maydis* ascospores.

**Supplementary Fig. 7: Macroscopic view of tar spot lesion.** (A) After the leaf has gone through senescence, stroma remain on the leaf surface, with a major vein running down the middle and the stromata forming around it. (B) Under a dissecting microscope, the clypeus and pycnidium are observed around the major vein. (C) Maize leaf with stroma. The blue arrow indicates the presence of a major vein.

