## Supplementary for "Uncovering the Infection Strategy of *Phyllachora maydis* during Maize Colonization: A Comprehensive Analysis"

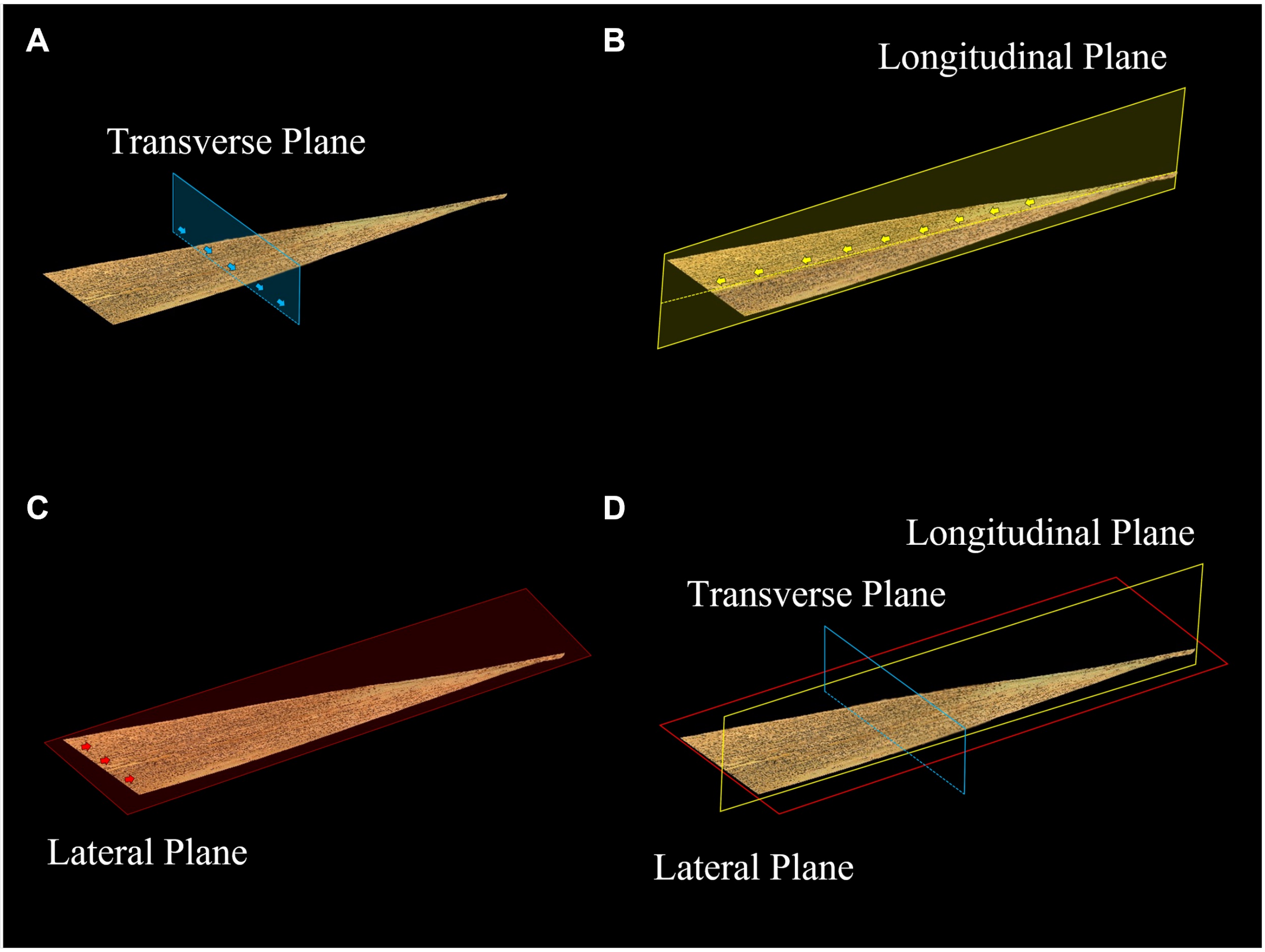


**Supplementary Fig. 1: Different orientation planes show the direction of cuts across the leaf blade.** (A) Transverse, or cutting the longitudinal axis at 90-degree angle. (B) Longitudinal, cutting along the long axis of the leaf blade perpendicular to the transverse plane. (C) Lateral plane, cuts the leaf horizontally top to the bottom, and (D) three planar cuts that together make a 3-demensional figure from 2-deminsional figures.

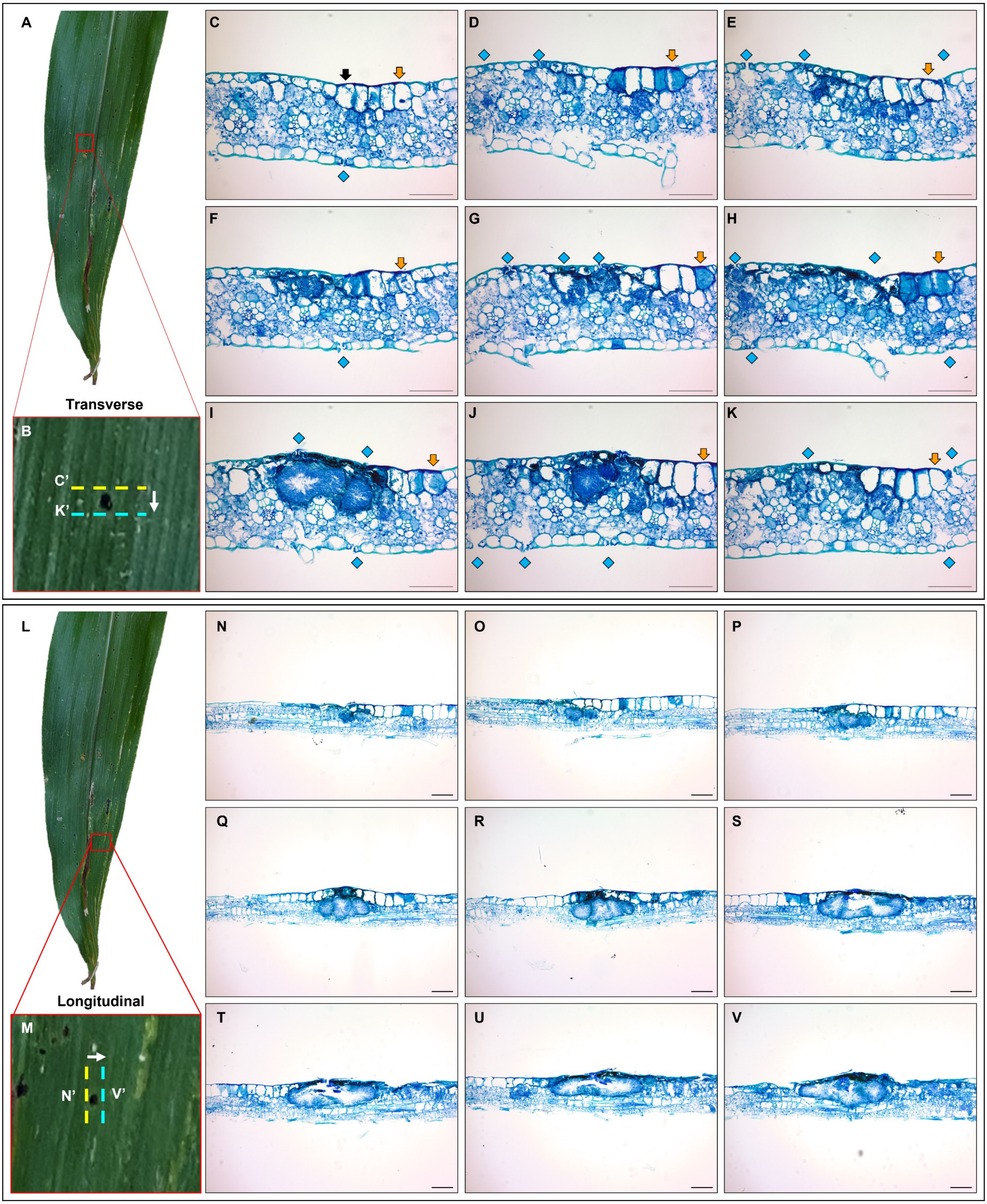

A
Transverse
B
C'
K'
C
D
E
F
G
H
I
J
K
L
Longitudinal
M
N'
V'
N
O
P
Q
R
S
T
U
V

**Supplementary Fig. 2: Transverse (A-K) and longitudinal (L-V) serial sections of stromata early in development.** (A) Leaf with stroma. (B) Section of leaf with stroma showing the direction of serial cuts (arrow). C' indicates the first cut; K' indicates the last. (C-K) Serial sections through one stromata. Because this is in the transverse plane, cell lines can be followed. The same cell line in each image (C-K) is indicated by an orange arrow. The blue diamonds are positioned over stomates, and the black arrow points to hyphae within a cell. Image C is on the outer periphery of the stroma, while Image I is in the center of the stromata. A well-developed clypeus is visible and immature pycnidia with conidiospores are developing. The young pycnidium is globose in shape, with a second lobe forming, and occupies the upper surface of the leaf while growing around the veins. (L) Leaf with stroma. (M) Section of leaf with stomata showing the direction of serial cuts (arrow). N' indicates the first cut; V' indicates the last. (N-V): Serial sections through one stromata. Hyphae in the upper epidermis form the clypeus and the side wall of the pycnidium emerges in deeper sections. By section Q, the pycnidium is no longer globose but has developed lobes. In the center of the stromata emerges (S-T), a long pycnidium with a clypeus, mature conidiospores, and a central ostiole are present. This long pycnidium remains on the upper side of the leaf and is outside the vasculature.

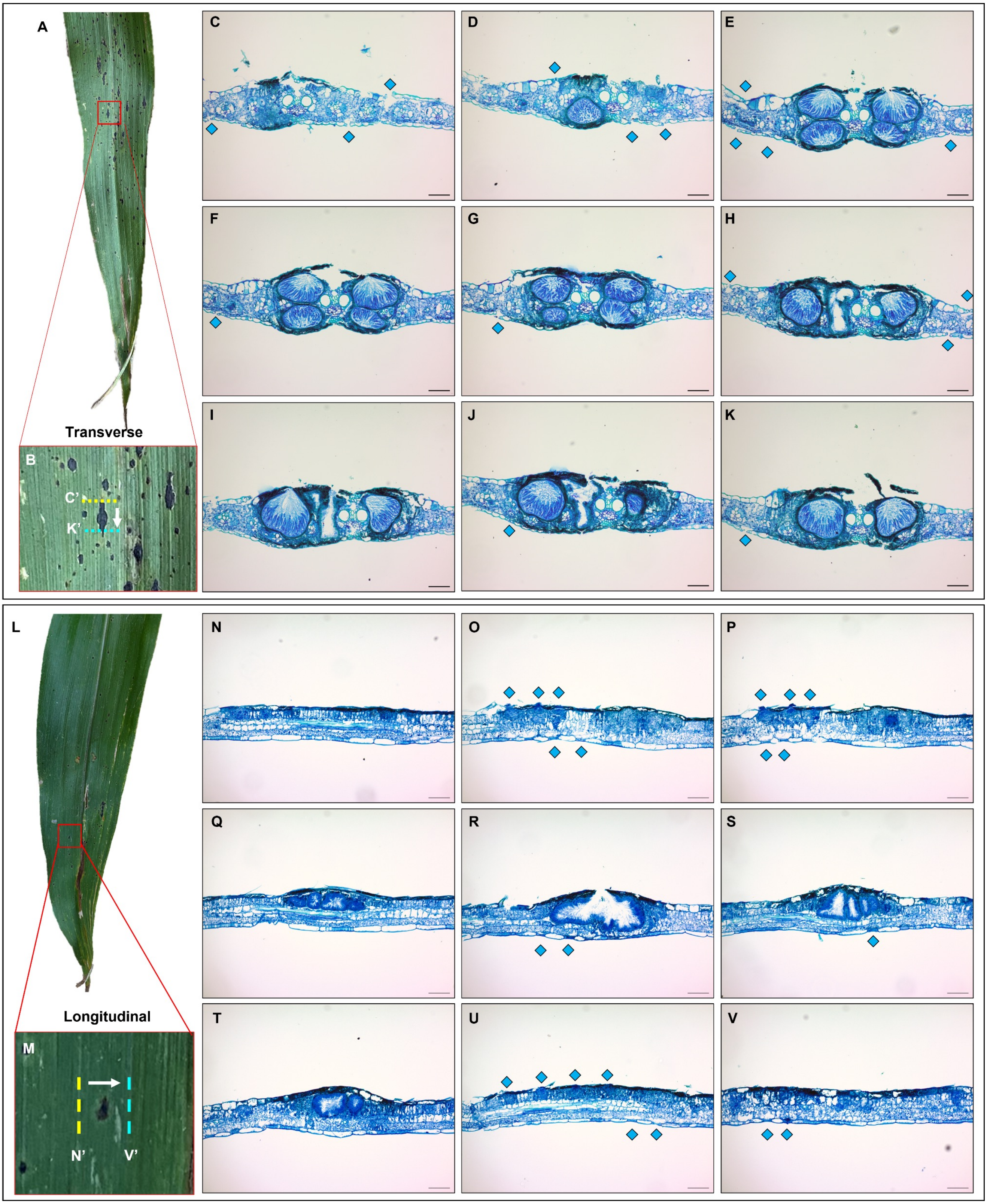
A
B
C'
K'
Transverse
C
D
E
F
G
H
I
J
K
L
M
N'
V'
Longitudinal
N
O
P
Q
R
S
T
U
V

**Supplementary Fig. 3**: **Transverse (A-K) and longitudinal (L-V) serial sections of stroma in mid-disease development.** (A) Leaf with stroma. In mid-development, the disease has spread to more of the leaf. The stoma has continued to develop and have increased more in the longitudinal than the transverse direction but do not overlap. (B) Section of leaf with stromata showing the direction of serial cuts (arrow). C' indicates the first cut; K' indicates the last. (C-K) Serial sections through one stromata. The blue diamonds are positioned over stomates. Sexual structures are present and an upper and lower clypeus in the epidermis. Moving through sections, perithecia surround the vasculature. The ostiole forms on the leaf abaxial side. (H-J) Closer to the center of the stromata, the pycnidium has developed. (L) Leaf with stroma. (M) Section of leaf with stomata showing the direction of serial cuts (arrow). N' indicates the first cut; V' indicates the last. The blue diamonds are positioned over stomates. (N-V): Serial sections through one stromata in the longitudinal orientation. (L-V), additional perithecia develop on the stromata periphery. The central pycnidium continues to develop multiple connecting lobes with a single opening through the adaxial clypeus. In longitudinal sections the increase in stromata length is apparent. This follows what is observed in the transverse sections: the disease travels along the cell lines with minimal transverse spread. Moving through serial sections reveals a developmental series of the perithecium, with the most mature closest to the pycnidium and the less mature at the outer edges of the stroma.

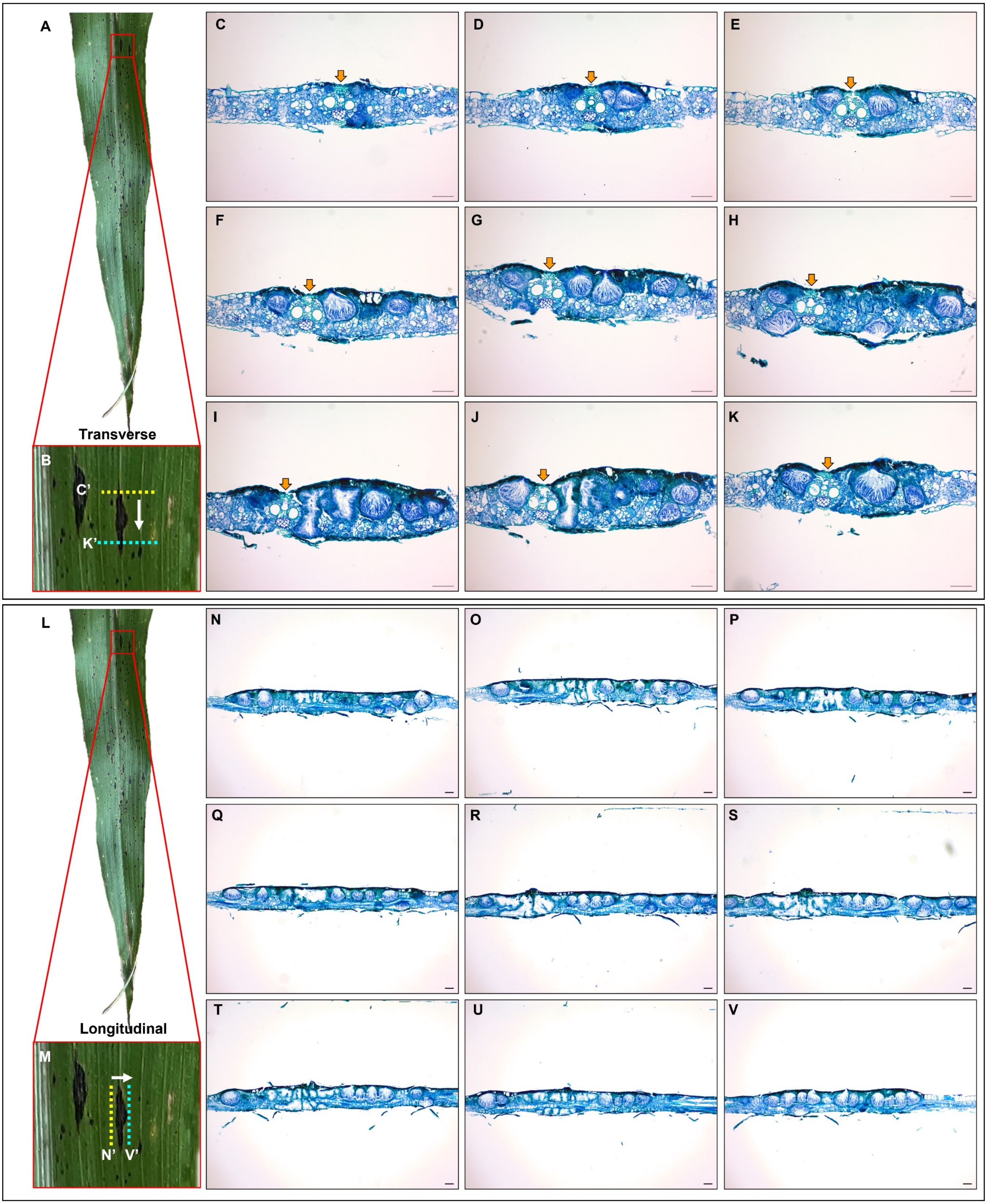

A
Transverse
B
C'
K'
C
D
E
F
G
H
I
J
K
L
Longitudinal
M
N'
V'
N
O
P
Q
R
S
T
U
V

**Supplementary Fig. 4: Transverse (A-K) and longitudinal (L-V) serial sections of stroma in late disease development.** (A) Leaf with stroma. In late development, the disease has spread throughout the leaf. The stroma has continued to develop and have increased more in the longitudinal than the transverse direction but do not overlap. (B) Section of leaf with stromata showing the direction of serial cuts (arrow). The disease follows the cell lines in the monocot leaf. C' indicates the first cut; K' indicates the last. (C-K) Serial sections through one stromata. The orange arrow points to the vasculature. Stroma do not form on top of each other, but smaller stroma form closer to larger stromata. An upper and lower clypeus stays directly over the reproductive structures. The vasculature is not occupied by *P. maydis*. The central multi-lobed pycnidium is present from one epidermis to the other (I-J). (L) Leaf with stroma. (M) Section of leaf with stomata showing the direction of serial cuts (arrow). N' indicates the first cut; V' indicates the last. (N-V): Serial sections through one stromata in the longitudinal orientation. The stromata has increased in length. In the center (S), a multi-lobed pycnidium is visible around the vasculature system (U). (S-V) Pycnidia are flanked by many perithecia.

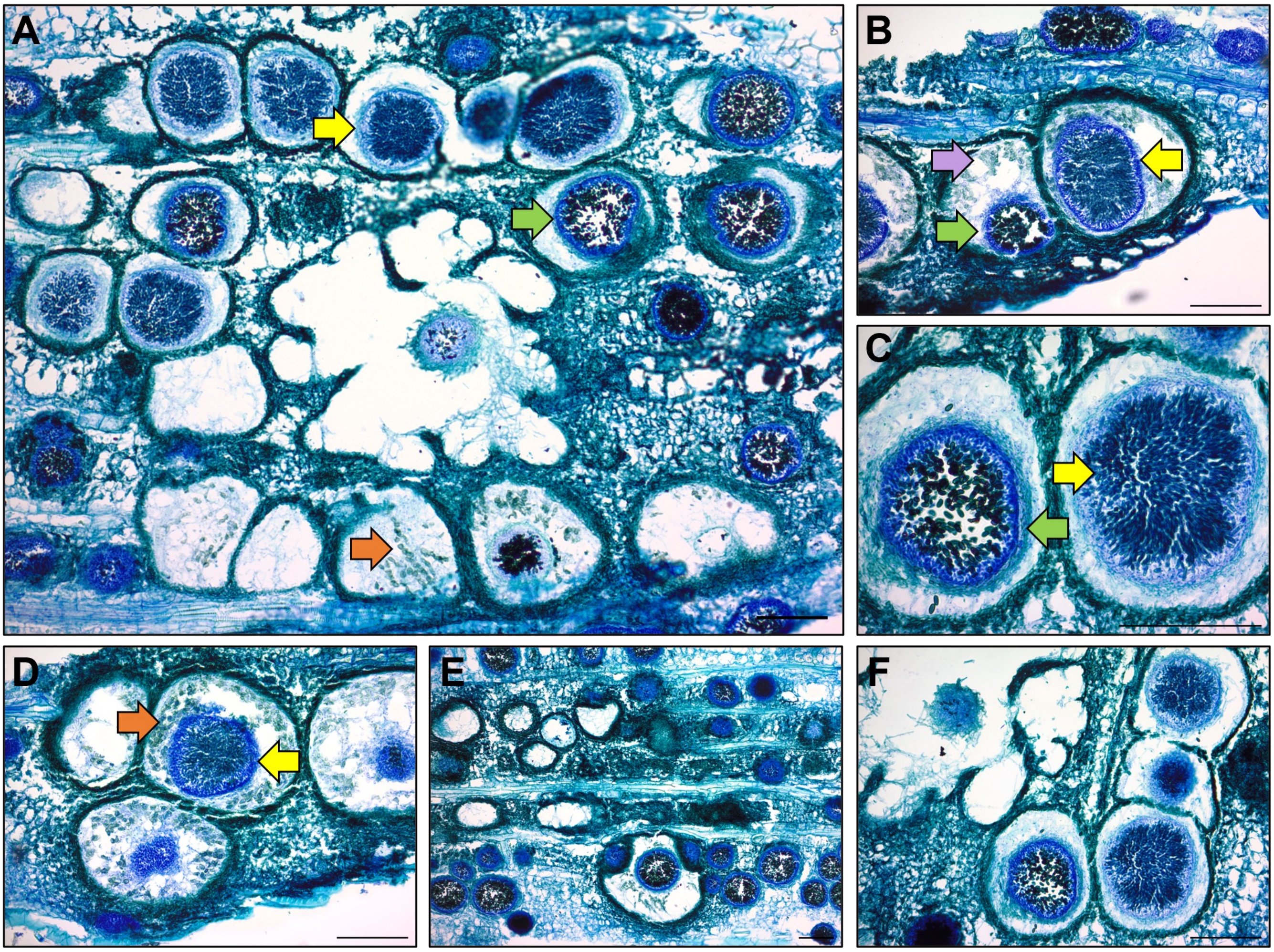


**Supplementary Fig. 5: *P. maydis* reproductive structures colonized by parasitic pathogens.** (A) Remnants of *P. maydis* pycnidium and perithecium with parasitic pycnidium inside. (B) *P. maydis* perithecia colonized by parasitic pathogens. The orange arrow indicates asci with ascospores still attached, the green arrow is suspected *Paraphaeosphaeria* spp., the yellow arrow indicates an unidentified parasite, purple arrow shows degrading *P. maydis* spores.

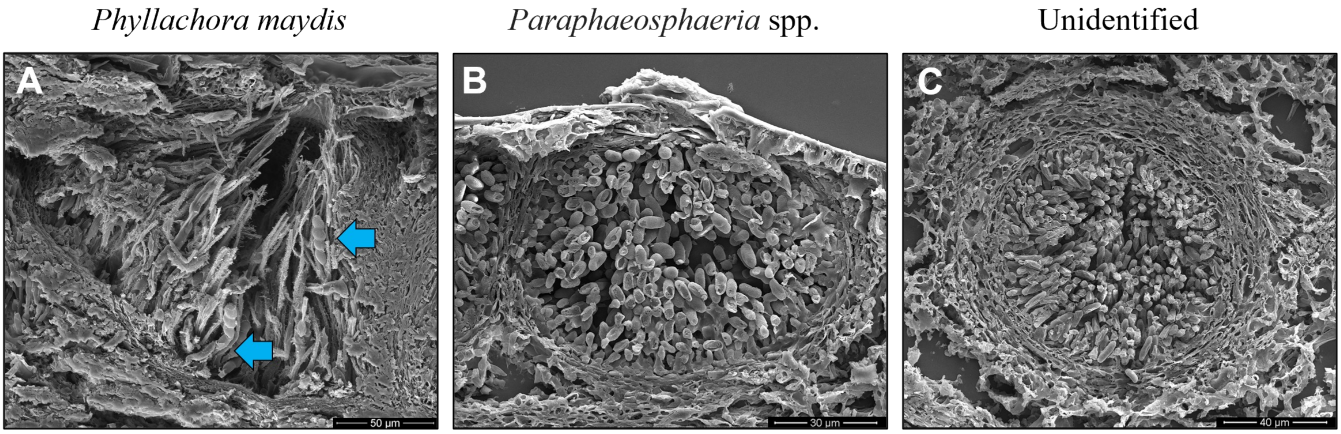


**Supplementary Fig. 6: SEM of different spore types found in fish eye lesions.** The blue arrows point to *P. maydis* ascospores.

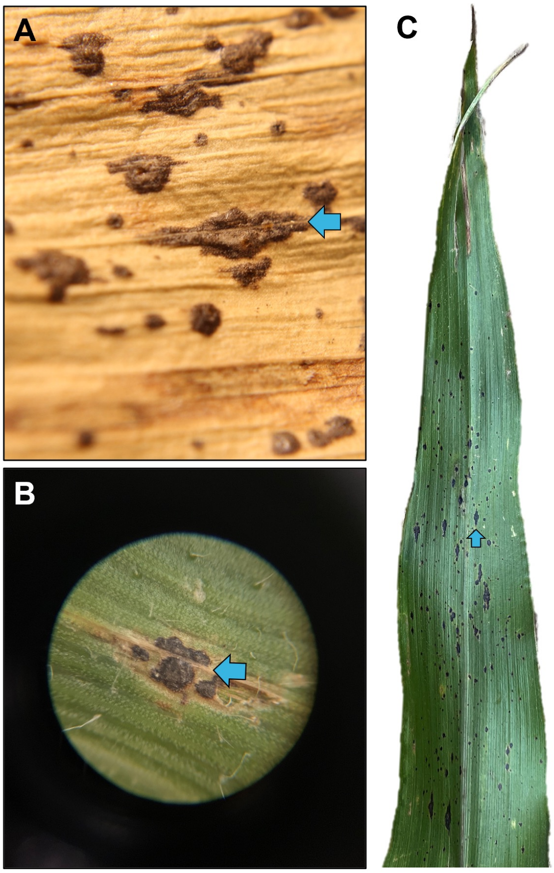


**Supplementary Fig. 7: Macroscopic view of tar spot lesion.** (A) After the leaf has gone through senescence, stroma remain on the leaf surface, with a major vein running down the middle and the stromata forming around it. (B) Under a dissecting microscope, the clypeus and pycnidium are observed around the major vein. (C) Maize leaf with stroma. The blue arrow indicates the presence of a major vein.
